# Lipid packing contributes to the confinement of caveolae to the plasma membrane

**DOI:** 10.1101/2025.03.13.643064

**Authors:** E. Larsson, A. Kabedev, H. Pace, J. Lindwall, F. Bano, J. Rae, R.G. Parton, C.A.S. Bergström, I. Parmryd, M. Bally, R. Lundmark

**Affiliations:** Department of Medical and Translational Biology and Laboratory for Molecular Infection Medicine Sweden, Umeå Centre for Microbial Research, SciLifeLab, Umeå University, 901 87, Umeå, Sweden; Department of Pharmacy, Uppsala University, Uppsala Biomedical Center, 751 23 Uppsala, Sweden; Department of Clinical Microbiology, Wallenberg Centre for Molecular Medicine & Umeå Centre for Microbial Research, Umeå University, 901 85, Umeå, Sweden; Institute for Molecular Bioscience, The University of Queensland, Brisbane, Queensland, Australia; Centre for Microscopy and Microanalysis, The University of Queensland, Brisbane, Queensland, Australia; Department of Medical Biochemistry and Cell Biology, University of Gothenburg, 405 30 Göteborg, Sweden

**Author notes:** Corresponding author: Richard Lundmark, Department of Medical and translational biology, Umeå University, 901 87 Umeå, Sweden.

**Keywords:** Caveolae, lipid packing, membrane curvature, internalization, lipid model membrane, molecular dynamics simulations, Dyngo-4a

## Abstract

Lipid packing is a fundamental characteristic of bilayer membranes. It affects all membrane-associated processes ranging from curvature generation to membrane fission. Yet, we lack detailed mechanistic understanding of how lipid packing directly affects these processes in cellular membranes. Here, we address this by focusing on caveolae, small 0-shaped invaginations of the plasma membrane which serve as key regulators of cellular lipid sorting and mechano-responses. In addition to caveolae coat proteins, the lipid membrane is a core component of caveolae that critically impacts both the biogenesis, morphology and stability of such membrane invaginations. We show that the small compound Dyngo-4a adsorbs and inserts into the membrane, resulting in a dramatic dynamin-independent inhibition of caveola dynamics. Analysis of model membranes in combination with molecular dynamics simulations revealed that a substantial amount of Dyngo-4a was inserted and positioned at the level of cholesterol in the bilayer affecting lipid order in a cholesterol dependent manner. Dyngo-4a-treatment resulted in decreased lipid packing of the plasma membrane. This prevented caveolae internalization and lateral diffusion without affecting their morphology, associated proteins, or the overall cell stiffness. Artificially increasing plasma membrane cholesterol levels was found to counteract the block in caveola dynamics caused by Dyngo-4a-mediated lipid packing frustration. Therefore, we propose that the outer leaflet lipid packing of cholesterol in the the plasma membrane critically contributes to the confinement of caveolae to the plasma membrane.

**Significance statement:** Larsson et al., demonstrate that lipid packing critically impacts the stability of small 0-shaped plasma membrane cavities termed caveolae. The plasma membrane (PM) release of caveolae is dynamin independent and halted by PM incorporation of the small compound Dyngo-4a, which leads to decreased lipid packing as characterized by the authors. The block in caveola internalization can be counteracted by increased PM levels of cholesterol which is enriched in caveolae and increases the lipid packing. This proof-of-principle study shows that lipid packing controls membrane budding generating membrane vesicles that remain stably associated with the membrane for an extended period of time.

## Introduction

The plasma membrane (PM) of eukaryotic cells has a distinctive asymmetric lipid composition with up to 40% cholesterol (chol) primarily located in the outer lipid leaflet (1). The PM lipid composition and asymmetry significantly influence lipid packing and membrane properties and thus also membrane remodelling processes such as the budding and scission of membrane vesicles (2, 3). During the formation of membrane vesicles, specific protein coats drive membrane curvature causing reorganization of lipids, asymmetry and changes in lipid packing (4). Classically, most attention has been directed towards vesicles such as clathrin coated vesicles (CCV), which rapidly undergo scission once the CCV has been formed and dynamin GTPases are recruited to the vesicular neck (4). Elucidation of the precise sequence of events leading to coated vesicle formation and scission have facilitated the generation of precise predictive models of vesicle formation (5). Other endocytic surface structures remain associated with the membrane for longer periods and their endocytic itineraries are far less well-defined (6, 7). Caveolae, which play important roles in lipid trafficking and monitoring of the PM composition and tension (8) show association with the plasma membrane and can stay connected for long time periods (9–11). Caveolae assemble by cooperative interaction of the integral membrane proteins caveolins 1-3 and peripherally attached cavins 1-4, which form the caveola coat (8, 12). The 0-shaped structure is stabilized by the ATPase EH domain containing 2 (EHD2), which multimerizes at the caveola (singular) neck thereby confining it to the PM (6).

Assembly of the caveola coat is highly dependent on distinct lipid species. In mammalian cells, caveolin 1 (Cav1) is thought to form homo-oligomers (8S complexes) in a chol dependent manner (13). The structure of the 8S complex was resolved using cryogenic electron microscopy and single-particle reconstruction, revealing a disc-shaped oligomere of 11 Cav1 subunits (14). Modelling of the lipid packing around the Cav1 oligomer showed that the Cav1 oligomer displaced the inner leaflet lipids, and that chol molecules together with sphingolipids formed a disordered monolayer in the outer leaflet facing the oligomer (15). At the PM, Cav1 8S-complexes assemble together with cavin1, generating membrane curvature resulting in typical 0-shaped caveolae (16, 17). Interestingly, flat caveolae structures positive for Cav1, cavin and EHD2 have been observed, suggesting that the curvature of the coat is adaptable (18). Removal of chol using Nystatin or methyl-β-cyclodexrin (MβCD) has been shown to result in flattening and disassembly of caveolae (19, 20) and release of cavins into the cytosol (21, 22). Along with chol, sphingomyelin (SM) and glycosphingolipids are also enriched in caveolae (14, 15, 23) and located in the outer lipid leaflet of the PM. The inner lipid leaflet of caveolae is enriched in phosphatidylserine (PS) and phosphatidylinositol 4,5 bisphosphate (PIP2) due to their interactions with cavins and EHD2 (23). In this way, lipid sorting by the caveolae coat and associated proteins generates a unique membrane domain with distinct lipid composition and high curvature.

While it is now apparent that a population of caveolae undergo scission to form endocytic carriers (9, 24–26) the regulation of their budding is still largely uncharacterized (24, 27, 28). Confinement to the PM is controlled by the ATPase EHD2. Loss of EHD2 results in more internalised caveolae, while overexpression or microinjection of EHD2 leads to very stable PM association (25, 26, 29). These effects have been quantified in live cells using TIRF microscopy by tracking the duration time of caveolin spots at the PM and by electron microscopy methods that distinguish budded from surface-connected caveolae (10, 11, 29, 30). The stabilizing effect of EHD2 is dependent on ATP-binding and the formation of ring-like oligomers of EHD2 at the caveolae neck (25, 30, 31). Although some caveolae remain PM associated for extended periods (24, 32), endocytosed caveolae can either re-fuse with the PM or fuse with endosomes while remaining as a distinct entity. The balance between surface association and internalization of caveolae is influenced by the lipid composition (29, 33). Increased PM levels of chol and glycosphingolipids has been found to increase curvature and promote caveola internalisation, while sphingomyelin prevents it. Yet, little is known about the mechanisms that control surface association and internalization. Recently, it was shown that caveola internalization is independent of dynamin (11), a GTPase which mediates scission of clathrin-coated pits from the PM (34). In fact, dynamin2 acted to prevent internalization of caveolae (11, 35).

Intriguingly, caveola internalisation has been shown to be inhibited by the small pyrimidyl-based compounds dynasore and Dyngo-4a resulting in more PM associated caveolae and less internal caveolae (9, 11, 21, 25). Dyngo is a modified version of dynasore with an additional hydroxyl group improving the activity (36). These compounds inhibit dynamin-mediated scission and the uptake of, for example, the transferrin receptor via CCVs (36, 37). Yet, cell viability and endocytosis via clathrin independent carriers (CLICs), which do not require dynamin for scission, is unaffected (36–39). Interestingly, Dyngo-4a constrains caveolae internalisation even in cells depleted of Dyn1 and Dyn2 (11), suggesting that the drug acts via another mechanism. Furthermore, both Dynasore and Dyngo-4a have been suggested to have wider effects on cells, for example affecting chol, lipid rafts, lamellipodia formation and cell migration (40, 41). Dynasore was shown to act protectively against chol-dependent pore forming toxins by disrupting PM lipid rafts (42). Dynamin depletion or inhibition using a dynamin inhibitor peptide showed that the effect was dynamin independent (42). Dyngo-4a-treatment has been shown to block membrane ruffling and inhibit fluid phase endocytosis in dynamin triple knockout (TKO) cells (43). These results clearly show that these compounds affect cellular processes at the plasma membrane that are dynamin independent and suggests that their actions are multifaceted. These insights have clouded the view on how this frequently used drug should be utilized in experiments addressing endocytosis and raise questions on how the resulting data should be interpreted.

Since Dyngo-4a-treatment dramatically affected caveola dynamics in Dyn2-depleted HeLa cells(11), we aimed to use this compound to reveal principles that control caveola dynamics. To obtain an understanding of the underlying mechanisms, we used interdisciplinary experimental setups and show that Dyngo-4a binds and inserts beneath the headgroups of the plasma membrane affecting the lipid order and membrane remodeling.

## Results

### The drug Dyngo-4a confines caveolae at the plasma membrane independently of dynamin

The dynamin inhibiting small molecules Dynasore and Dyngo-4a (Fig. S1A) have been demonstrated to prevent caveolae internalization (11, 21, 25). However, Dyngo-4a was shown to have a dramatic stabilizing effect on caveola dynamics even in cells depleted of Dyn2 (11). This suggested that the drug acts via an unknown mechanism, which, if clarified, might advance our understanding of the principles that control caveola dynamics. To assure that the effect of Dyngo-4a was totally independent of all dynamin isoforms we utilized the well-established dynamin TKO cell line where the addition of tamoxifen can be used to silence the three different isoforms of *dnm* (43). This abolishes dynamin protein expression compared to the untreated control (ctrl) cells (Fig. 1A) (43). Analysis of the caveolae core proteins Cav1, cavin1 and EHD2 by fluorescence microscopy showed that, similarly to control cells, typical punctate caveola structures covered the PM of TKO cells (Fig. 1B and Fig. S1B-C). Quantification of the density of caveola based on Cav1 punctae revealed no significant difference between the wild type (wt) and knock out cells (Fig. 1C), in agreement with previous results (18).

**Figure 1.**
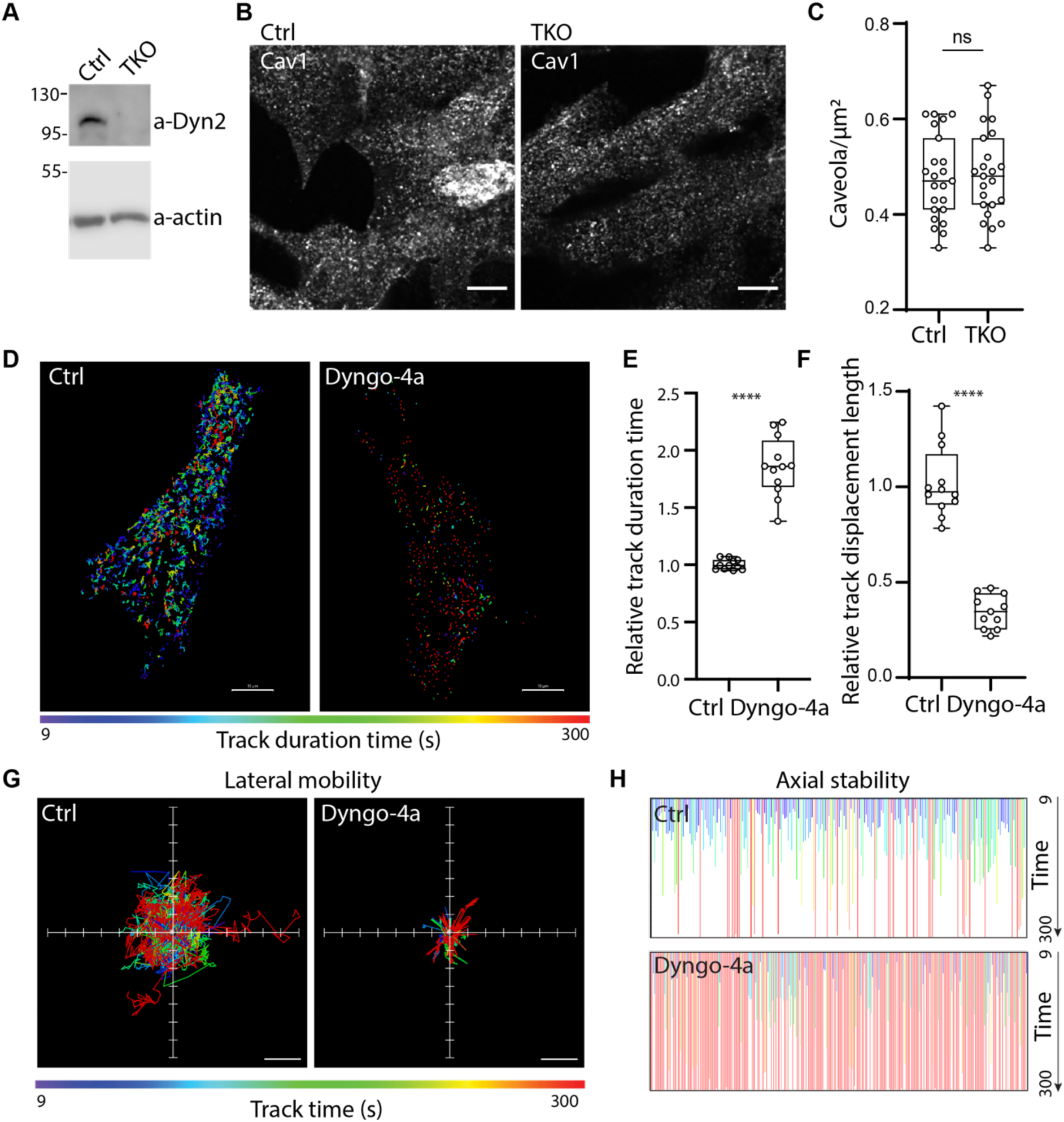
The drug Dyngo-4a constrains both axial and lateral mobility of caveolae independently of dynamin. **(A)** Immunoblots of dynamin triple knock out cells showing dynamin 2 expression after 5 days of Tamoxifen-treatment (KO) or no treatment (Ctrl) as indicated. **(B)** Representative immunofluorescent staining of Caveolin1 in dyn TKO cells treated as in (A) as indicated, Scale bar, 5 μm. **(C)** Quantification of the number of caveolae per square micrometer counted in the basal PM in immunofluorescent labeled dyn TKO cells treated as in (A) as indicated. Mean ± SD from at least 20 cells per condition. Significance was assessed using *t* test, p =0.7421. **(D)** Color-coded trajectories showing caveolae movements in the PM. Cav1-GFP was transiently expressed in dyn TKO cells after 5 days of Tamoxifen treatment. Cells were serum-starved 1 h prior to 30 min treatment with DMSO (Ctrl) or 30 μM Dyngo-4a before imaged on TIRF over 5 min. Red trajectories represent caveolae with long duration times and purple represents caveolae with short duration times as indicated. Scale bar, 10 μm. **(E-F)** Quantification of Cav1-GFP track duration time (E) and track displacement length (F). Numbers were related to ctrl-treated cells, and track mean from at least 23 cells per condition are shown ± SD (E) and track length mean from at least 15 cells per condition are shown ± SD (F). Significance was assessed using t test, ****p ≤ 0.0001. **(G)** Visualization of the track displacement of color-coded trajectories in (D) by alignment of the starting position of all tracks, representing lateral mobility. **(H)** Kymographic visualization of the track duration of color-coded trajectories in (D) that were present at time zero, representing axial stability.

To monitor the dynamics of caveolae, Cav1-GFP was transiently expressed in Tamoxifen-treated TKO cells. Cells were treated for 30 min with either Dyngo-4a or DMSO and Cav1-GFP low-expressing cells were then imaged using total internal reflection fluorescence (TIRF) microscopy to visualize Cav1-GFP at, or in close proximity to the basal PM (Video 1 and 2). Using single particle tracking, where Cav1-GFP punctae of a specific size and fluorescent intensity were identified, we could determine their lateral mobility (displacement length) and axial movements in and out of the TIRF field (duration time) (Fig. 1D-F and Fig. S1F) (10, 11, 29, 30). Tracking of a large number of Cav1-GFP puncta from both ctrl and Dyngo-4a-treated cells, showed that the average duration times were significantly increased and that the displacement length was greatly reduced in the Dyngo-4a-treated cells as compared to ctrl-treated cells (Fig. 1E and F). Detailed analysis of the lateral mobility of caveolae showed that both directed transport and Brownian motion decreased even in tracks with long duration time, following Dyngo-4a treatment (Fig. 1F and G). This is indicative of fewer released and mobile caveolae as well as less membrane undulations (11). Furthermore, in Dyngo-4a-treated cells, the majority of the caveolae present at time zero were tracked persistently throughout the movie (Fig. 1H). In fact, almost no short duration tracks were detected (Fig. 1G) which would be expected if Dyngo-4a treatment resulted in progressive loss of Cav1-GFP punctae from the PM. To validate that the Cav1-GFP punctae were caveolae and not disassembled caveolin structures, Dyngo-4a treated cells were fixed and stained against endogenous cavin1 and EHD2. This showed that Dyngo-4a treatment did not disrupt the colocalization of these caveolae markers to Cav1-GFP structures as compared to ctrl treated cells (Fig. S1D and E). In addition, when mCherry-cavin1 was co-expressed the Cav1-GFP punctae were consistently positive for cavin1 during Dyngo-4a treatment (Video 3). This showed that Dyngo-4a treatment did not lead to caveola fragmentation and disassembly of the caveola complex.

### Dyngo-4a does not affect caveola morphology but stalls their internalization

To test if caveolae maintained their typical morphology and make sure that caveolae did not flatten in response to Dyngo-4a treatment, we analyzed the PM of A431 cells in thin sections using electron microscopy (Fig. 2A). To account for differences in caveola size caused by random sectioning, we measured the width, height and neck diameter of individual surface connected caveola. By comparing the ratios of height and neck to width we found no significant difference in terms of neck size, width or height of caveolae in Dyngo-4a treated cells as compared to ctrl cells (Fig. 2B). These data showed that caveolae displayed the characteristic 0-shape following Dyngo-4a treatment.

**Figure 2.**
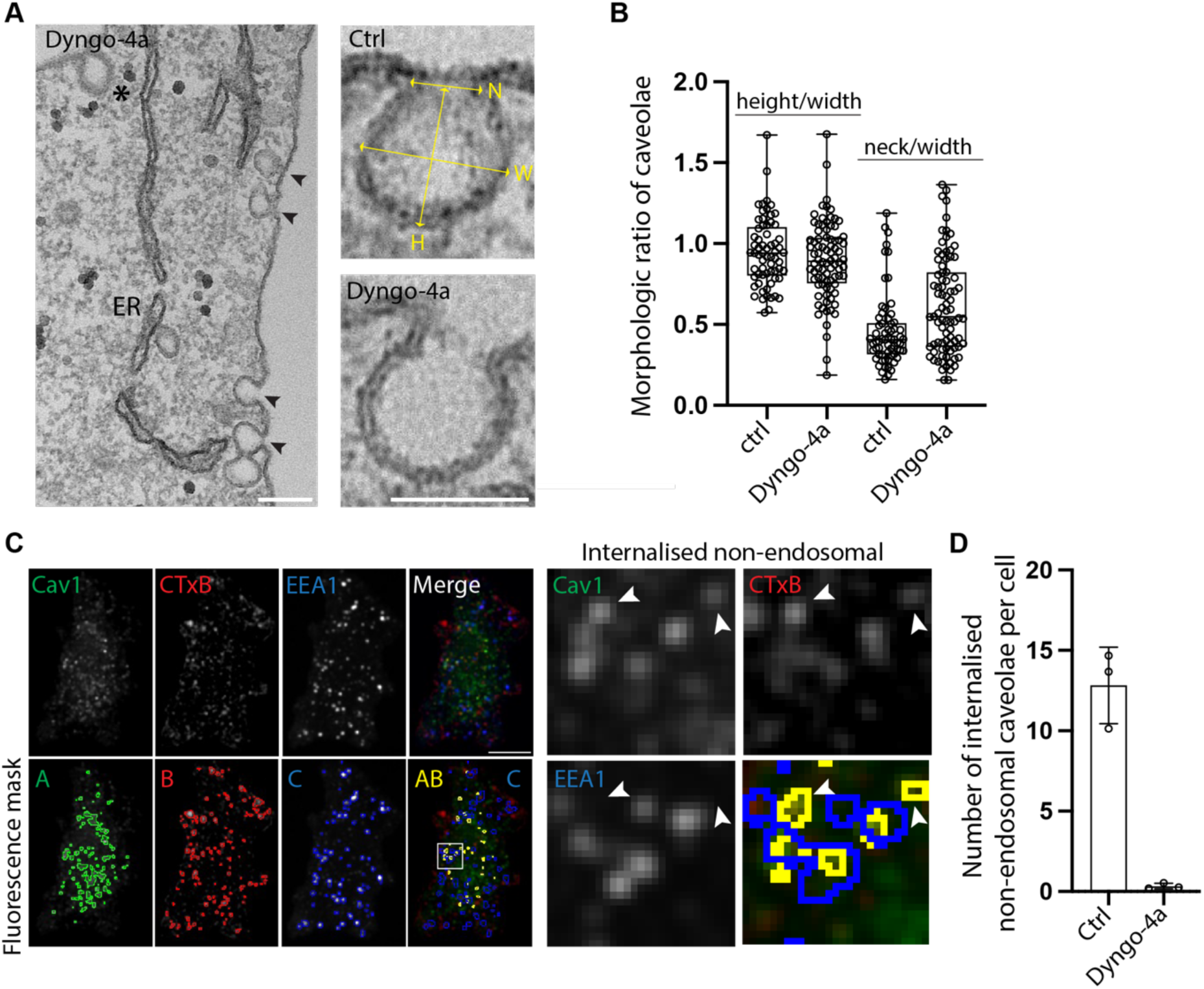
Caveolae internalization is inhibited by Dyngo-4a. **(A)** Representative electron micrographs of caveolae in Dyngo-4a or ctrl treated cells. Black arrowheads show caveolae and stars show CCV in cells treated with Dyngo-4a. Yellow arrows mark the positions where neck (N), width (W) and height (H) measurements were acquired. Scale bar, 100 nm. **(B)** Scatter plots show the ratios of height or neck to width measurements of 60 caveola per treatment, mean ± SD. **(C)** Representative immunofluorescent maximum-intensity projections of immunofluorescent CTxB uptake in ctrl-treated cells. Bottom panels show the masks generated from each fluorescent marker which were used to quantify the number of Cav1-GFP and CTxB-647 colocalizing spots devoid of EEA1, indicated by the white arrows in the magnified view. **(D)** Quantification of the number of Cav1-GFP/CTxB colocalizing spots negative for EEA1 per cell in ctrl or Dyngo-4a treated cells. Mean ± SD from 15 cells per condition (n=3). Significance was assessed using t test, *p ≤ 0.05.

Although the decreased duration time observed in our live cell TIRF data following Dyngo-4a treatment suggest that internalization of caveolae is reduced, this is not directly measuring internalization. Quantification of caveolae endocytosis is challenging since there is no known cargo specifically endocytosed by caveolae, and that internalized caveolae are not easily differentiated from other vesicles based on morphology in EM. Therefore, to verify that the internalization of caveolae was affected, we used Alexa-647 labelled cholera toxin B subunit (CTxB) to mark internalized caveolae in Cav1-GFP expressing cells (Ref). Since CTxB is also endocytosed by other pathways, confocal microscopy and image analysis of whole cells was used to specifically identify caveolae (Cav1-GFP spots) positive for CTxB (Fig 2C and S2A). To discriminate internalized caveolae from vesicles that already had fused and delivered CTxB to early endosomes, cells were additionally stained against the early endosomal marker EEA1 (Fig. 2C and S2A). Quantification of caveolae that were CTxB-positive and EEA1 negative revealed a striking effect on the number of internalized caveolae in Dyngo-4a treated cells as compared to ctrl cells (Fig. 2D). This showed that Dyngo-4a prevent caveolae endocytosis. To further scrutinize this effect, we photobleached a region of the cell and used confocal imaging to monitor the recovery of Cav1-GFP punctae in the cytosol (Fig S1G and H). When analyzed, cells treated with Dyngo-4a had lower fluorescence recovery rate than ctrl-treated cells showing that less cytosolic Cav1-GFP puncta were present following Dyngo-4a treatment (Fig. S1H). Taken together, this demonstrated that Dyngo-4a-treatment prevents release of caveolae from the PM.

### Dyngo-4a adsorbs and inserts into the lipid bilayer

Because Dyngo-4a is an amphiphilic compound (Fig. S1A) we reasoned that it might interact with the lipid bilayer and influence the properties of the membrane. To decipher if Dyngo-4a inserts into membranes, we created a lipid monolayer of 1-palmitoyl-2-oleoyl-glycero-3-phosphocholine (POPC) lipids suspended at an air-water interphase and monitored the changes in the lateral pressure after injection of the compound using Langmuir-Blodgett trough tensiometry (Fig. 3A, illustration)(44). Injection of Dyngo-4a under the equilibrated lipid monolayer resulted in a steep rise of the surface pressure (Fig. 3A). These results show that Dyngo-4a indeed inserts between the lipids of the planar monolayer. Injection of a control solution containing the same DMSO concentration as the Dyngo-4a solution, had a minor effect on the surface pressure (Fig. 3A), in agreement with the known interaction between DMSO lipid bilayers (45).

**Figure 3.**
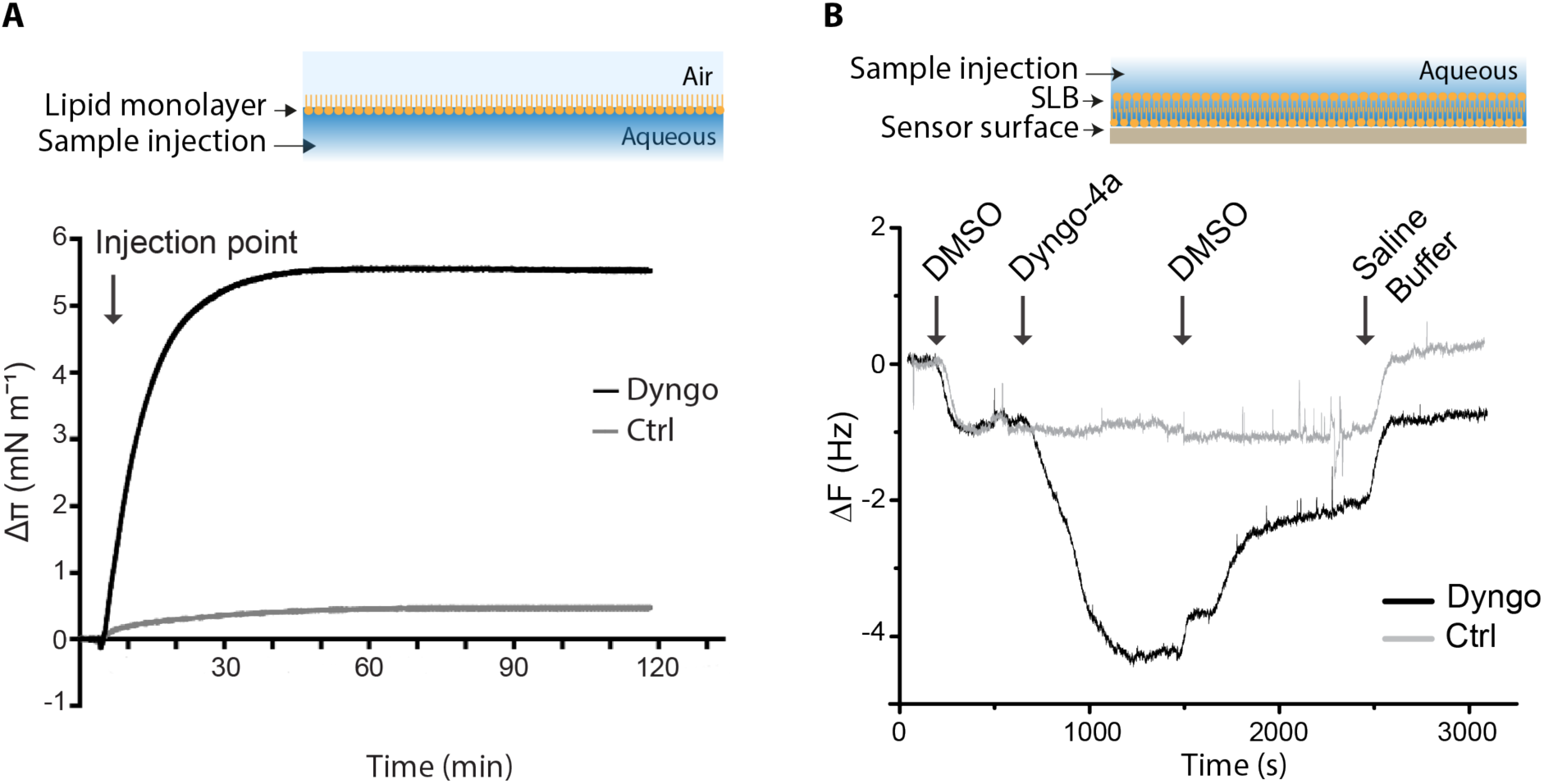
Dyngo-4a binds and inserts into membranes. **(A)** Top, Illustration of Langmuir-Blodget experiment. Bottom, representative Dyngo-4a adsorption to POPC monolayer. Dyngo-4a (30 μM) (black line) or DMSO (gray line) was injected underneath the lipid film with the starting surface pressure of 20 mN m^-1^and the surface pressure shift (Δπ) was recorded over time. **(B)** Top, illustration of the QCM-D set up. Bottom, QCM-D measurements of SLB formation and Dyngo-4a (30 μM) (black line) or DMSO (gray line) adsorption to a POPC SLB, followed by rinsing. Arrows depict time of injection of DMSO (equivalent to the concentration that Dyngo-4a is dissolved in), sample injection (Dyngo-4a or DMSO), DMSO injection and saline buffer wash (150 mM NaCl, pH 7.4) injection respectively.

To further confirm the surface interaction between Dyngo-4a and the membrane, we used quartz crystal microbalance with dissipation monitoring (QCM-D). This technique uses an acoustic sensor to track adsorption and desorption to the sensor surface, which is expressed as a function of frequency shift (τ1*f*) (Fig. 3B, illustration)(46). Supported lipid bilayers (SLBs) of POPC were generated on the sensor surface via vesicle fusion. The system was first equilibrated in buffer containing DMSO. Upon addition of Dyngo-4a in the same buffer the frequency decreased substantially, indicating that Dyngo-4a adsorbed to the SLB (Fig. 3B). Rinsing the SLB with buffer led to desorption of Dyngo-4a, showing that binding was at least partially reversible in DMSO-containing buffer (Fig. 3B). After returning to DMSO-free buffer, the system was of low enough softness (Fig. S3), in order to use the Sauerbrey equation to convert the frequency change into adsorbed mass. A comparison of the calculated mass of the SLB alone and the calculated mass of the Dyngo-4a-containing-SLB allowed us to estimate the amount of Dyngo-4a retained after washing (0.9 ± 0.3 Hz), accounting for 6 ± 2 mol% (Fig. 3B).

### Dyngo-4a inserts at a similar position as cholesterol decreasing the lipid order in MD simulations

To gain detailed information about how Dyngo-4a is positioned in the membrane, we performed molecular dynamic (MD) simulations where Dyngo-4a was added to a POPC bilayer. The free sampling simulations were performed for 200 ns and showed that Dyngo-4a was incorporated into the bilayer underneath the headgroups (Fig. 4A). This is in agreement with the hydrophobic properties of the compound. To gain information about the energy landscape with barriers and preferred regions of positioning of Dyngo-4a within the membrane, Dyngo-4a was sampled through the POPC membrane using simulations with a convolved adaptive biasing potential based on the Accelerated Weight Histogram (AWH) method. The potential of mean force (PMF) profile showed that Dyngo-4a preferred the area of the POPC membrane, around 1.1 to 1.2 nm from the bilayer center (Fig. 4B). Interestingly, an energy difference of approximately 40 kJ/mol between the lowest PMF point and the center of the membrane (Fig. 4B) suggests that Dyngo-4a is unlikely to cross the POPC bilayer spontaneously.

**Figure 4.**
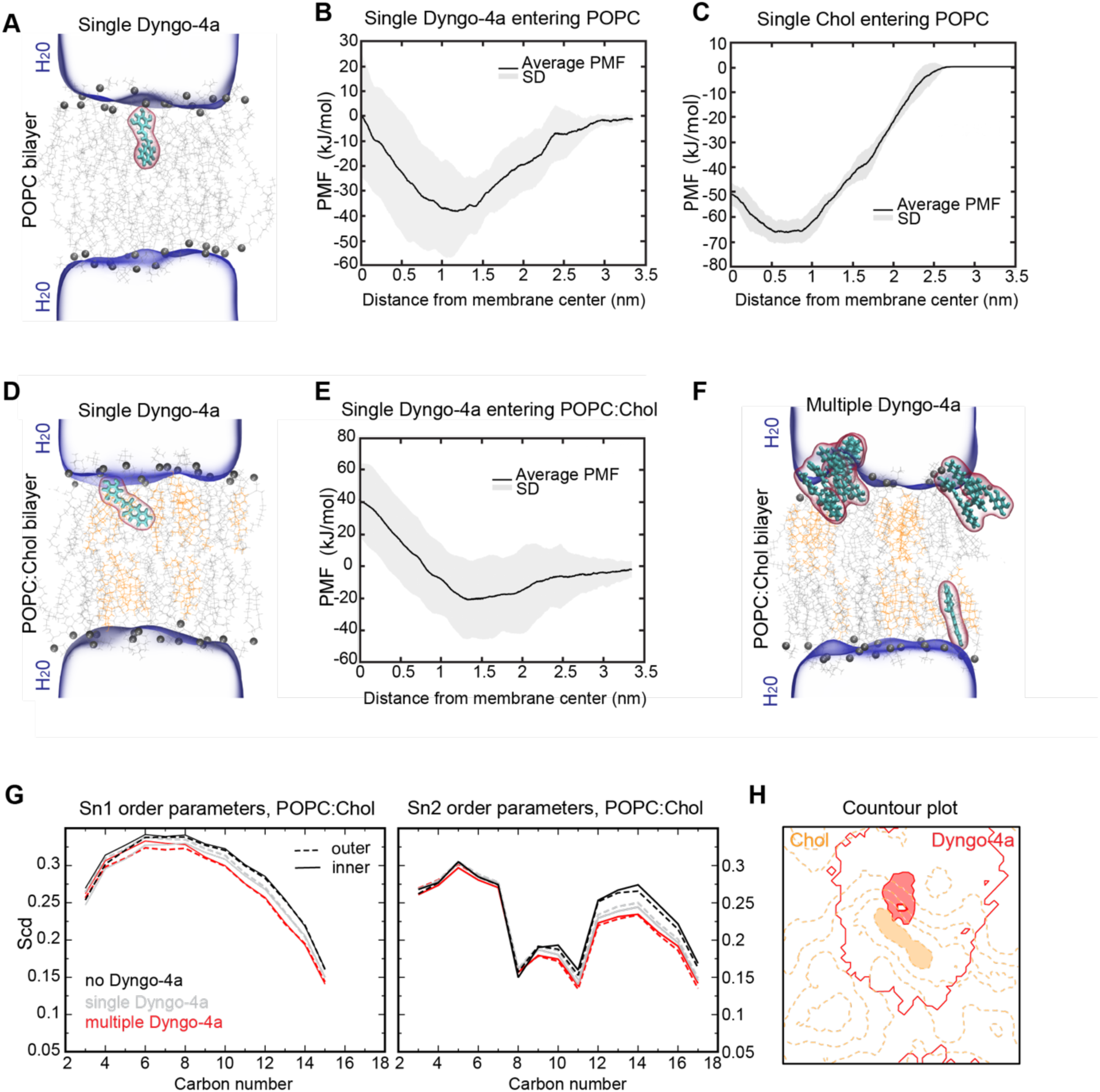
Dyngo-4a inserts underneath the phospholipid headgroups at a similar position as cholesterol. **(A)** Molecular dynamic simulation of Dyngo-4a in a membrane. Snapshot of a single Dyngo-4A molecule (cyan stick and red volume) in POPC bilayer (phospholipid headgroups in dark gray and lipid chains as light gray sticks, dark blue represents the aqueous phase) **(B-C)** AWH simulations of Dyngo-4a (B) or Chol (C) in POPC membrane. PMF profile show the energy required to move a Dyngo-4a molecule or a Chol molecule from the membrane center to the aqueous phase, respectively. **(D)** Molecular dynamic simulation of Dyngo-4a in a membrane. Snapshot of a single Dyngo-4A molecule (cyan stick and red volume) in POPC:Chol (70:30%) bilayer (phospholipid headgroups in dark gray and lipid chains as light gray sticks, chol as orange sticks, dark blue represents water). **(E)** AWH simulations of Dyngo-4a in POPC:Chol (70:30%) membrane. The PMF profile shows the energy required to move a Dyngo-4a molecule from the membrane center to the aqueous phase. In the PMF profiles the lines represent the mean and the shaded regions represent the SD from triplicate simulations. **(F)** Molecular dynamic simulation snapshot of multiple Dyngo-4a molecules in POPC:Chol (70:30%) bilayer (color coding as in (D)). Dyngo-4a molecules did not cross the membrane throughout the simulation so they were placed randomly at the beginning of the simulations and could translocate from one leaflet to another via periodic boundary condition. **(G)** Deuterium order parameter profiles for POPC tails in POPC:Chol (70:30%) membrane. Left show sn1 tail (saturated, 16 carbons) and right shows sn2 tail (unsaturated, 18 carbons). No Dyngo-4a (black), single Dyngo-4a (gray), multiple Dyngo-4a (red), dashed lines depict the outer leaflet and solid line the inner leaflet. **(H)** Contour plot based on heatmaps in Fig. S3E highlighting the predominant configuration whit Dyngo-4a (red) localized adjacent to a chol (orange) cluster in the outer leaflet.

As chol is highly abundant within the PM, we hypothesized that Dyngo-4a may interact with or displace chol within the lipid tail region due to partial spatial overlap. AWH-enhanced simulations of a chol molecule interacting with the POPC membrane showed that the center of mass located in regions (0.5 to 1.0 nm) overlaps with the preferred region of Dyngo-4a (Fig. 4C and B). To test if addition of chol to the membrane would alter the position of Dyngo-4a, MD simulations were performed on a membrane consisting of 70% POPC and 30% chol. This showed that the chol molecules localized to the POPC tail region as expected. In this setup Dyngo-4a was still incorporated under the headgroups (Fig. 4D). However, AWH simulations showed that in the presence of chol, the preferred Dyngo-4a position shifted slightly closer to the head group region and exhibited a broader distribution compared to pure POPC (Fig. 4E). This was further supported by the density profiles of Dyngo-4a overlaid with POPC and chol (Fig. S4A). Furthermore, the mid-membrane energy barrier that Dyngo-4a would need to overcome to pass the membrane was increased to approximately 60 kJ/mol (Fig. 4E). Notably, when multiple Dyngo-4a molecules were added to the limited membrane area of the simulation single molecules inserted in the bilayer. However, within the simulation time most of the molecules aggregated and formed stacks that were not able to entirely penetrate through the headgroup region of the membrane (Fig. 4F and Fig. S4B). These aggregates nonetheless perturbed the membrane structure, particularly in the POPC:Chol membrane, as confirmed by order parameter measurements (Fig. 4G and Fig. S4C). The presence of chol increased the order parameter, whereas the addition of a single Dyngo-4a or multiple molecules generally lowered the order of the lipids by up to 0.02 and 0.04 units, respectively (Fig. 4G and Fig. S4C). Both leaflets were affected, with a slightly greater effect in the outer leaflet. Alignment of the POPC sn1 tails were disrupted throughout the tail length, while deviations in the alignment of the sn2 tails were more pronounced in the distal carbons farthest from the headgroups (Fig. 4G). In pure POPC membrane the effect of both single molecules and clusters were less pronounced (Fig. S4C). Concurrently, when the average lateral space each lipid occupies within the membrane plane was measured, the area per lipid increased in the mixed POPC:Chol membranes upon insertion or adsorption of Dyngo-4a (Fig. S4D), whereas it remained largely unchanged in POPC membranes. Finally, colocalization analysis revealed that Dyngo-4a preferentially associated with chol clusters in the membrane (Fig. 4H and Fig. S4E), contrasting with its broader distribution in pure POPC and with the chol density profile in the absence of Dyngo-4a. These results further support that Dyngo-4a insertion significantly alters lipid packing in POPC:Chol membranes. Moreover, while chol increases lipid order, Dyngo-4a has the opposite effect.

### Membrane insertion of Dyngo-4a decreases lipid packing in the plasma membrane

Next, we aimed to address how membrane incorporation of Dyngo-4a affected the properties of the PM. Interestingly, we found by analyzing TIRF time-lapse images that Dyngo-4a majorly impaired the typical undulations of PM protrusion and retraction in a concentration-dependent manner (Fig. 5A-B and S5 A-B). This resulted in a “frozen” cell membrane with no membrane ruffling and lamellipodia formation which has been reported previously as a dynamin-independent effect of Dyngo-4a (43). Similarly, we observed a steady and continuous concentration-dependent decrease in lateral mobility of caveolae as concentrations of Dyngo-4a were increased (Fig. S5C). Quantification of the effect on lateral movement over time revealed a fast initial decrease in response to Dyngo-4a and that the maximum level of inhibition was reached after 20 to 25 minutes (Fig. S5D). This is comparable to effects on clathrin-coated vesicle (CCV) endocytosis as measured by the internalization of transferrin (Tfn), although the inhibition of this process was slightly faster and completely blocked already at 10 μM of Dyngo-4a (Fig. S5C). Taken together, these data showed that Dyngo-4a affects several active processes that involve remodeling of the PM.

**Figure 5.**
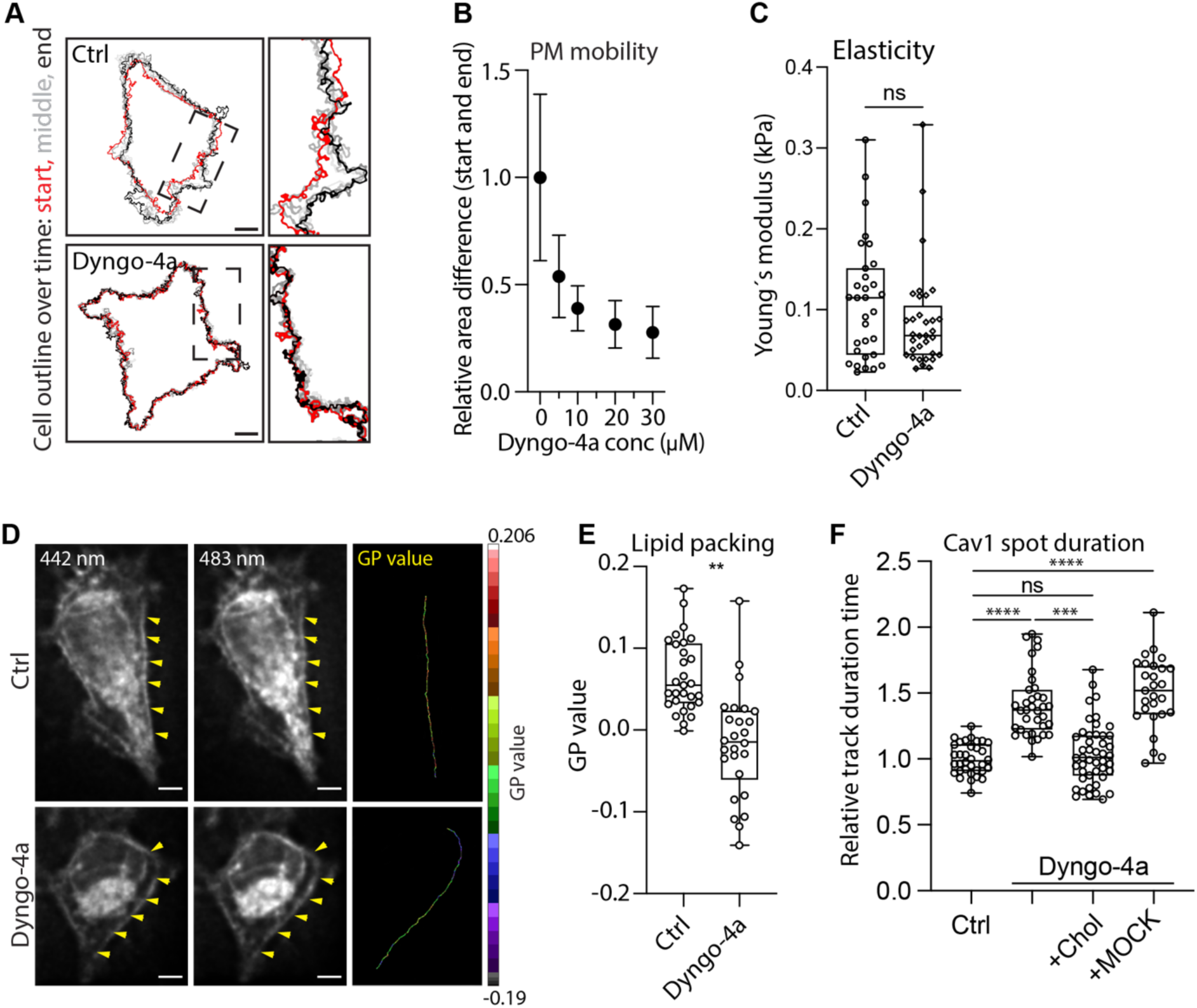
Dyngo-4a decreases the lipid packing in the outer leaflet of the PM which is counteracted by increased levels of cholesterol. **(A)** Visualization of the PM mobility of a ctrl-treated cell (top) and a Dyngo-4a-treated cell (bottom) from 5 minute TIRF movies. The outline of the PM at the start (0 s, red), middle (60, 120, 180 and 240 s, gray) and end (300 s, black) are depicted. Insets to the right show magnifications of the indicated areas. Scale bar, 10 μm. **(B)** Quantification of the PM mobility of cells following treatment with DMSO (ctrl) or Dyngo-4a at different concentrations as indicated. The means is shown from five cells per condition ± SD. **(C)** Young moduli calculated from force-indentation curves from the representative curves in (Fig S4) over 31 and 33 cells treated with DMSO and Dyngo-4a, respectively. Each data point is an average value calculated from at least 40 individual force curves on each cell. Statistical analysis: parametric student t-test with Welch’s correction. ns = non significant. **(D)** Representative deconvolved fluorescent micrographs of HeLa cells stained with C-Laurdan captured at 442 nm (left panel) and 483 nm (middle panel). Yellow arrows depict analyzed stretch of PM with no neighboring fluorescent internal membrane as seen in the right panel. Red area corresponds to membrane with high membrane order and purple corresponds to membrane with low membrane order. Scale bar, 5 μm. **(E)** Quantification of the mean GP-values for DMSO (ctrl) or Dyngo-4a treated cells as indicated. All data points from at least 26 cells per condition are shown, n = 3, mean ± SD. Significance was assessed using *t* test, ** p ≤ 0.0065. **(F)** Quantification of Cav1-mCh track duration time following chol:MβCD treatment as indicated. Numbers were related to ctrl-treated cells. All data points from at least 26 cells per condition are shown, n = 3, mean ± SD. Significance was assessed using t test, ****p ≤ 0.0001.

To determine if Dyngo-4a affected the mechanical properties of the membrane, we quantified Young’s modulus (a measure of the elastic properties of materials) using an atomic force microscopy (AFM) -based nanomechanical setup. Young’s moduli for ctrl cells and cells treated with Dyngo-4a were obtained by fitting force-indentation curves with the Hertz model (Fig. S5F). Quantification revealed that Dyngo-4a treatment had no significant effect on the elastic modulus, showing that the drug does not affect the overall cell stiffness (Fig. 5C). The simulation data showed that membrane insertion of Dyngo-4a influenced the lipid packing by decreasing the lipid order. To analyze if Dyngo-4a affected this property of the plasma membrane bilayer, we used the commonly employed fluorescent probe 6-dodecanoyl-2-[*N*-methyl-*N*-(carboxymethyl)amino] naphthalene (C-Laurdan) to measure the degree of lipid packing in the PM (47). Simulations on C-Laurdan suggests that it does not flip in between lipid leaflets and therefore this probe reports on the outer leaflet lipid packing when added to the extracellular media (48). The fluorochrome of the probe inserts into the interphase between the lipid headgroups and the acyl chains of the lipid bilayer, and emit different wavelengths depending on the level of water penetration in the membrane. First, we confirmed that Dyngo-4a had a similar effect on caveolae duration in the presence or absence of C-Laurdan (Fig. S5E). C-Laurdan was added to cells treated with Dyngo-4a and z-stacks were acquired using fluorescent microscopy. Using the intensities from the different emission channels, the relative membrane order was established by calculating the generalized polarization (GP) value of regions of the PM (Fig. 5D). Quantification of the GP-values showed that the PM of Dyngo-4a treated cells displayed a significantly lower mean GP-value than ctrl-treated cells (Fig. 5E), showing that the compound decreases the lipid packing in agreement with the simulation data. Based on this, we propose that membrane insertion of Dyngo-4a increases the hydrated hydrophilic area in the outer lipid leaflet, causing lipid packing frustration.

### Increased PM levels of cholesterol counteracts the inhibition in caveola dynamics caused by Dyngo-4a

Our simulation data showed that the positioning of Dyngo-4a in the lipid leaflet was similar to chol and that Dyngo-4a preferentially associated with chol clusters and affected the spatial distribution of chol in the membrane. Therefore, we hypothesized that Dyngo-4a could affect chol in the outer leaflet of the PM leading to decreased lipid packing which might be the cause of halted caveolae internalization. To test if increased levels of chol would restore caveolae dynamics, we used fusogenic liposomes to incorporate chol in the PM (29). However, Dyngo-4a treatment prevented fusion of the fusogenic liposomes (Fig S5F). This suggests that the decreased lipid packing mediated by Dyngo-4a also affects membrane fusion. Addition of chol prebound to MβCD to cells has previously been reported to successfully increase the PM-levels of chol (49). Using this approach with fluorescently labelled chol, we could visualize efficient incorporation of chol using TIRF microscopy (Fig S5G). This methodology allowed us to analyse how addition of MβCD-chol to Dyngo-4a-treated cells affected the caveola duration time. Strikingly, increasing the chol levels following Dyngo-4a treatment resulted in a caveolae track duration time similar to ctrl cells (Fig 5F). This rescue effect was not observed in cells where Dyngo-4a treatment was followed by incubation in media alone. These data showed that the block in caveola dynamics induced by Dyngo-4a could be counteracted by increased levels of chol in the PM.

Taken together, our data show that membrane incorporation of Dyngo-4a results in decreased lipid packing and PM confinement of caveolae, which can be overcome by increasing the PM levels of chol. Based on this we conclude that lipid packing is a critical determinant in the balance between internalization and stable PM association of membrane vesicles.

## Discussion

Lipid composition and membrane properties are known to be critical determinants of both caveolae biogenesis and endocytosis (50). Here, we show that the small pyramidyl-based compound, Dyngo-4a, inserts into membranes affecting lipid packing and resulting in a dramatic, dynamin-independent, effect on caveola internalization. This enabled the use of this compound to identify lipid packing as a driving force behind confinement of such highly curved invaginations to the PM. Detailed characterization of the membrane interaction of Dyngo-4a using model membranes showed that the compound was inserted into a lipid monolayer. Simulations employing AWH showed that single Dyngo-4a molecules were positioned just beneath the headgroup region of the lipids. Interestingly this coincides with the preferred localization of chol and indeed the positioning of Dyngo-4a was influenced by the presence of chol. Furthermore, we observed chol-dependent membrane adsorption and shallow insertion of Dyngo-4a aggregates. Both types of insertion contributed to decreased lipid order, especially in the outer leaflet, in the simulations. Using SLBs and the QCM-D system we could determine that a substantial amount (6 ± 2 mol%) of Dyngo-4a was indeed adsorbed to the bilayer after washing. Taking this value as a reference, the amount of Dyngo-4a bound to the bilayer before rinsing, which resulted in a frequency shift of 3 Hz, was extrapolated to be about 15.6 mol%. Yet, as indicated by the simulations both inserted Dyngo-4a and adsorbed aggregates would contribute to this amount. Therefore, it is difficult to precisely determine the amount of Dyngo-4a inserted into cellular membranes.

Analysis of the PM of cells treated with Dyngo-4a using C-Laurdan revealed an increased water accessibility of the outer leaflet of the membrane as compared to ctrl cells. This showed that incorporation of the compound caused lipid packing frustration increasing the hydrated hydrophilic area, which will influence the properties of the membrane. We could not detect any significant effect on the overall stiffness of the cells implying that the decreased packing does not influence the general deformability of the PM. Furthermore, highly curved invaginated caveolae and CCVs were still present in cells as confirmed by EM, and previous results show that clathrin independent endocytic carriers are actively formed in Dyngo-4a-treated cells (38). This suggests that Dyngo-4a incorporation does not have a generic impact on the elasticity of the membrane. However, membrane insertion of Dyngo-4a increased the average duration time of caveolae detected at the PM and reduced the number of internalized caveolae. Dyngo-4a-treated cells were almost completely devoid of short tracks and no scission events could be detected. In fact, we found that internalization of CTxB via caveolae decreased by more than 95 percent. In addition, track analysis also showed that short tracks typically derived from membrane undulations interfering with the TIRF field were absent. This is in agreement with the finding that PM movements were stalled in Dyngo-4a treated cells (43). Therefore, we conclude that the decreased lipid packing caused by Dyngo-4a prevents both the internalization of caveolae as well as membrane undulations. The effect that lipid packing has on caveolae internalization is in line with the model suggesting that the caveola coat proteins together with specific lipids drives scission as a continuum of membrane curvature generation (29). Lipid packing around the caveolin 8S complex was recently proposed to drive membrane curvature by inducing elastic stresses of tilt and splay in the monolayer proximal to the caveolin complex (51). Furthermore, recent simulations of the lipid packing around the caveolin 8S complex show that the lipids are displaced by the Cav1 oligomer in the inner leaflet. In the outer leaflet facing the complex, 40-70 chol molecules and sphingolipids form a disordered monolayer (15, 52). The differential contact energy between the hydrophobic phase of the disc and the outer membrane lipids versus between outer and inner leaflet lipids favors the accumulation of chol on the 8S discs. This provides molecular insight into how lipids are sorted by the 8S complex and how lipid packing might control the curvature, where increased chol concentration in the PM would lead to more tilt and splay and membrane stress, and eventually scission of caveolae (reviewed in (53)). Our data suggests that Dyngo-4a is restricted to the outer lipid bilayer in a similar position as chol. Thus, incorporation of Dyngo-4a might be expected to interfere with lipid packing of the disordered monolayer facing the Cav1 oligomer preventing increased curvature and scission. Interestingly, although not statistically verified, we observed more caveolae with broader neck diameters Dyngo-4a treated cells. Indeed, our data show that artificially increased levels of chol in the PM can counteract the block in caveola internalization induced by Dyngo-4a. These data further support outer leaflet lipid packing as a potential critical determinant of membrane scission.

Based on previous literature and our data, Dyngo-4a efficiently blocks endocytosis of both CCVs and caveolae. We and others have also shown that some clathrin-independent endocytic pathways are still operative under conditions in which Dyngo-4a blocks clathrin and caveolar endocytosis (36, 38). This demonstrates that, despite the general effects on membrane properties reported here, there is considerable specificity in terms of regulation of specific cellular processes. Therefore, the numerous studies that have used this drug to block endocytosis of caveola and CCVs in relation to other endocytic pathways are still valid. Yet, according to our data, the inhibition of caveolae internalization is independent of dynamin and instead due to the insertion of Dyngo-4a in the outer lipid bilayer and thereby decreasing the lipid packing. Currently, we cannot address if this effect is specific to caveolae, or general to the different on- and off-targets effects that has been ascribed to this drug. Dyngo-4a has not been shown to pass membranes and enter cells and our data show that the energy barrier for Dyngo-4a to pass from the outer to the inner leaflet is high. This would imply that in order for Dyngo-4a to directly interact with dynamin in the cytosol, membrane passage would have to be mediated by, for example, a transporter. Yet, it is difficult to definitively discern if Dyngo-4a acts via one or several inhibitory mechanisms. Our comparison of the effects of Dyngo-4a treatment on CCV endocytosis and caveola mobility suggests that CCV endocytosis is more sensitive to Dyngo-4a in terms of time and concentration. However, these differences could either be due to Dyngo-4a acting via more than one mechanism or the same mechanism of action having quantitatively different impacts on CCVs and caveolae due to their different sensitivities to perturbations of specific properties of the membrane.

Our results show that caution should be taken when conclusions are drawn based on the use of small inhibitory compounds. The drug Pitstop 2, developed to inhibit clathrin assembly and thus clathrin mediated endocytosis has also been reported to freeze the membrane (54). Currently, very few small compounds used for chemical screening are tested for their ability to insert into membranes. Yet, we show Dyngo-4a can be used to efficiently block endocytosis of both CCV and caveola, and that such compounds can be very useful when addressing how membrane properties affect cellular processes. Based on this work, we propose that lipid packing is instrumental for the balancing of lateral movement and internalization of caveolae.

## Acknowledgment

We acknowledge the Biochemical Imaging Center (BICU) at Umeå University within the National Microscopy Infrastructure, NMI (VR-RFI 2016-00968) and the Microscopy Australia Research Facility at the Center for Microscopy and Microanalysis at the University of Queensland for providing support and assistance with microscopy. We especially thank Irene Martinez at BICU for assistance and expertise with image analysis and data visualization. We acknowledge the Biochemical Macromolecular Characterization Umeå at Umeå university (BMCU). The computations were enabled by resources provided by the National Academic Infrastructure for Supercomputing in Sweden (NAISS), partially funded by the Swedish Research Council through grant agreement no. 2022-06725. We thank Jeremy Adler for support with the script for C-Laurdan software analysis and James Rae for assistance with electron microscopy. We thank Chloe Williams for assistance with the RMSD calculations. The work was supported by grants to RL from the Swedish Research Council (dnr 2021-05117) the Swedish Cancer Society (CAN 23 3004 Pj 01 H) and grants to IP from Magnus Bergvall’s Foundation. RGP was supported by an Australian Research Council (ARC) Laureate Fellowship (FL210100107). HP and FB were supported by the Knut and Alice Wallenberg foundation.

## Author contributions

Elin Larsson and Richard Lundmark designed the research and Elin Larsson performed and analysed the molecular biology, live cell imaging and tracking experiments. Aleksei Kabedev performed modelling experiments, Jakob Lindwall tracking experiments, Hudson Pace QCM-D experiments and analysis, Fouzia Bano AFM experiments and analysis, Ingela Parmryd enabled the C-Laurdan measurements and performed their analysis. Robert Parton and James Rae enabled the electron microscopy experiments and performed their analysis. Christel A. Bergström and Marta Bally provided financial contribution. Elin Larsson and Richard Lundmark conceived the work and wrote the manuscript, and all authors contributed with text, figures and editing.

## Materials and methods

### Reagents

Dyngo-4a (ab120689, Abcam) was dissolved in DMSO to make a stock solution of 30 mM. For experiments the working concentration was 30 µM unless otherwise stated and Ctrl samples were treated with equivalent DMSO concentration (0.1%) unless otherwise stated. 4-hydroxy-tamoxifen (Sigma) was dissolved in ethanol (99%) to make a stock solution of 10 mM. C-Laurdan was a generous gift from Dr. Bong Rae Cho, Korea University. Human Transferrin Alexa Fluor 647-conjugate (T23366) was bought from Invitrogen. POPC (1-palmitoyl-2-oleoyl-*sn*-glycero-3-phosphocholine) was purchased from Avanti Polar Lipids Inc (Alabaster). DMSO (34943), chloroform (CHCl_3_), and methanol (MeOH) were purchased from Sigma-Aldrich.

### Cell lines and constructs

The dynamin triple knockout cells were a kind gift from professor De Camilli (43). The cells were maintained in DMEM supplemented with 10% (vol/vol) FBS and 1x penicillin-streptomycin. The HeLa Flp-In T-REx Caveolin1-mCherry cells were previously generated by us (29). Cells were maintained in DMEM supplemented with 10% (vol/vol) FBS, 100 μg/ml hygromycin B (Thermo Fisher Scientific), and 5 μg/ml blasticidin S HCl (Thermo Fisher Scientific) for plasmid selection. Expression at near endogenous levels was induced by incubation with 0.5 ng/ml doxycycline hyclate (Dox; Sigma-Aldrich) for 16 to 24 h prior to imaging. All cell lines were cultivated at 37°C, 5% CO_2_ in a humidified incubator and tested negative for mycoplasma.

### Cell treatments

Ablation of *dynamin 1-3* from Dynamin TKO cells was achieved by culturing the cells with 2 μM 4-hydroxy-tamoxifen for two days. Cells were split to prevent overcrowding and cultured with 300 nM 4-hydroxy-tamoxifen for an additional three days prior to experiment. Protein levels were analyzed by SDS-PAGE and immunoblotting to Amersham Hybond P 0.45 polyvinylidene difluoride membrane (MERCK) using rabbit anti-Dyn2 (PA1-661; Thermo Fisher Scientific), and mouse anti-β-actin (3700; Cell Signaling). Following HRP-conjugated secondary antibodies were used: goat anti-rabbit (AS09 602; Agrisera) and goat anti-mouse (A9917; MERCK). Dynamin TKO cells were transfected with Lipofectamine 2000 (Thermo Fisher Scientific) using Opti-MEM I reduced serum medium (Thermo Fisher Scientific) for transient protein expression of Cav1-GFP according to manufacturer’s instructions. For Dyngo-4a treatment, cells were serum-starved 1 hour prior to addition of 30 μM Dyngo-4a in live cell medium for 30 min prior to experiment or as indicated in Fig. 4 B and D-E. DMSO (0.1%) was used as control. For C-Laurdan experiments, Cav1-mCherry HeLa FlpIn TRex cells were incubated in 3μM C-Laurdan in imaging buffer (phenol-red free DMEM) containing catalase and glucose oxidase (7000 units/ml and 16 units/ml, respectively) at 37°C, 15 min prior to imaging. Fusogenic liposomes were prepared from a lipid mixture of DOPE, DOTAP, and Bodipy-tagged cholesterol at a ratio of 47.5:47.5:5. Lipid blends were in MeOH:CHCl_3_ (1:3, vol/vol). Following the generation of a thin film using a stream of nitrogen gas, the vesicles were formed by addition of 20 mM Hepes (VWR, pH 7.5, final lipid concentration 2.8 μmol/ml) and incubated for 1.5 h at room temperature. Glass beads were added to facilitate rehydration. The liposome dispersion was sonicated for 30 min (Transsonic T 310, Elma). Ctrl or Dyngo-4a treated cells were incubated with fusogenic liposomes (7nmol/ml) and incorporation of the fluorescent lipid in the PM was imaged using TIRF microscopy. For chol: MβCD -complex treatment, MβCD was dissolved in serum free media (phenol-red free DMEM) supplemented with HEPES (25mM) to give a concentration of 5 mM. Chol in CHCl_3_ was added to a glass vial and a thin film of chol was created using a stream of nitrogen gas while rotating the vial on an angle. Thereafter, nitrogen gas was flowed into the vial for an hour to evaoporate remaining solvent. The chol was resuspended by vortexing in freshly made MβCD solution (5mM) for a final chol concentration of 480 mg/ml. The chol: MβCD solution was sonicated for 10 x 5 sec and incubated at 37°C ON with constant stirring. The chol:MβCD mixture was diluted 1:1 and cells were incubated with the solution and imaged using TIRF microscopy. For visualization of the incorporation, chol and Bodipy-chol at a ratio of 90:10 was complexed to MβCD and used as described above.

### Immunostaining and transferrin and CTxB uptake

Tamoxifen-treated and non-treated (ctrl) dynamin TKO cells were seeded on high precision coverslips (No. 1.5H, Paul Marienfeld GmbH and Co. KG) in 24-well plates at 3× 10^4^ cells/well and incubated overnight (37°C, 5% CO_2_). Cells were fixed with 3% PFA in PBS (Electron Microscopy Sciences) for 15 min at room temperature (RT) and subsequent permeabilization and blocking was carried out simultaneously using PBS containing 5% goat serum and 0.05% saponin. Cells were then immunostained with rabbit anti-Caveolin1, rabbit anti-PTRF (RRID:AB_88224) both from Abcam, rabbit anti-EHD2, RRID:AB_2833022 (25), followed by goat anti-rabbit IgG secondary antibody coupled to Alexa Fluor 488, (Thermo Fisher Scientific) as previously described (55). Co-staining of caveolae components was not technically possible due to that mouse antibodies did not work in the mice cell line. Confocal images were acquired using a Zeiss Spinning Disk Confocal microscope controlled by ZEN interface (RRID:SCR_013672) with an Axio Observer.Z1 inverted microscope equipped with an EMCCD camera iXonUltra from ANDOR. For Transferrin uptake, Cav1-mCherry HeLa FlpIn TRex cells were seeded on precision coverslips in 24-well plates at 6 × 10^4^ cells/well and induced with Dox and incubated at 37°C, 5% CO_2_ 16-18 hours prior to uptake assay. Cells were pretreated with Dyngo-4a or DMSO as specified in Fig. 4 D-E and then incubated 10 min at 37°C, 5% CO_2_ with Alexa flour-647 conjugated transferrin (Thermo Scientific) (5 μg/ml). Cells were washed with ice-cold PBS and kept on ice for a total of 15 min and immediately fixed with 3% PFA for 15 min at RT. Images were aquired using a Leica DMI8 inverted Thunder widefield microscope equipped with a LEICA DFC 9000 GTC camera controlled by Leica Application Suite X 3.9.1.28433. For CTxB uptake, cells were seeded in 24-well plates and prepared in the same way as described above prior to uptake. Alexa-647 CTxB (Thermo Fisher Scientific) (1 μg/ml) was added to cells and incubated at 37°C, 5% CO_2_ for 5 minutes. Cells were then washed with ice-cold PBS 3 times 5 minutes and immediately fixed with 3% PFA for 15 min at RT. Immunofluorescence staining against mouse-anti-EEA1 (BD Biosciences) followed by goat anti-mouse IgG secondary antibody coupled to Alexa Fluor 568, (Thermo Fisher Scientific) was executed as described above. Confocal Z-stack images were acquired using a Zeiss Spinning Disk Confocal microscope controlled by ZEN interface (RRID:SCR_013672) with an Axio Observer.Z1 inverted microscope equipped with an EMCCD camera iXonUltra from ANDOR. Micrographs were prepared using Fiji, RRID:SCR_002285 (56) and Adobe Photoshop v25.6.

### Live cell fluorescence microscopy imaging

Dynamin TKO cells were treated with 4-hydroxy-tamoxifen for five days in total and the day before experiments the cells were seeded on glass coverslips (CS-25R17, Warner instruments) in 6-well plates at 1.5 × 10^5^ cells/well and incubated at 37°C, 5% CO_2_. T-REx cells were induced with 0.5 ng/ml Dox and seeded on glass coverslips (CS-25R17) in 6-well plates at 3 × 10^5^ cells/well and incubated at 37°C, 5% CO_2_. Before imaging, the media was replaced with live-cell media (DMEM high glucose, no phenol red (Gibco), supplemented with 1 mM sodium pyruvate (Gibco) and imaged at 37°C with 5% CO_2_. For TIRF-microscopy, images were acquired for 5 min at 3 s intervals (every third image with definitive focus) using a Zeiss Axio Observer.Z1 inverted microscope that was equipped with an EMCCD camera iXon Ultra from ANDOR and an alpha Plan-Apochromat TIRF 100×/1.46 oil objective controlled by ZEN software. Only the basal membrane was analyzed as the laser penetration depth can not be adjusted to image the apical and lateral plasma membrane. All micrographs and videos were prepared with Fiji (56) and Adobe Photoshop CS6. C-Laurdan imaging was performed on a wide-field fluorescence Nikon Ti-U microscope (Nikon, Tokyo, Japan) equipped with a Cascade 1 k camera and a dual viewer (Photometrics, Tucson, AZ). A 60X water objective (NA 1.27) and refractive index-matched immersion oil (Zeiss, Oberkochen, Germany) were used. Excitation was with a LED at 380 nm (CoolLED, Andover, UK). A FF01-387/485/559/649 excitation filter (Semrock, Lake Forrest, IL) and a z365/577rpc dichroic (Chroma, Rockingham, VT) were used in the microscope and in the dual viewer a ms-470LDX dichroic (Chroma) and emissions filters 442/46 and FF01–483/32 from Semrock were used. Stacks of eleven images with a z-step size of 200 nm were acquired close to the basal membrane of the cells.

### Analysis of caveola dynamics

Imaris software 10.0.1 (Bitplane) was used for tracking analysis of Cav1-GFP or Cav1-mCh positive structures segmented as spots as previously described (29). Structures with a diameter of 0.4 μm were tracked and the applied algorithm was based on Brownian motion with max distance traveled of 0.8 μm and a max gap size of 3. Statistics for track duration and track displacement length from three independent experiments were extracted and analyzed. Tracks in figures were displayed using the spectrum colormap for track duration showing the caveolae heterogeneity within a cell where red tracks represent stable caveolae and blue/purple highly dynamic caveolae. The number of quantified cells is specified in the respective figure legend. Unpaired two-sample *t* test was performed on track duration (s) and track displacement (μm) using Prism 10.0.2, and data are shown as fold change.

### Analysis of PM dynamics

Masks of the total area of the basal membranes of the first and last image of 5 minute TIRF movies were established based on the membrane epiflouroscense using ImageJ. To quantify the PM motion, the area change was calculated by subtracting the end mask from the start mask (1) and by subtracting the start mask from the end mask (2) using the Image J image calculator. The total area change was calculated by subtracting the result from (1) and (2) from the area of the start image. The number of quantified cell areas are specified in respective figure legend. Unpaired two-sample *t* test was performed and data are shown as fold change. The root mean square deviation (RMSD) was computed using sections of membrane profiles extracted from control and Dyngo-4a (30uM) treated cells with gridded overlays. Start and end frame of five min time-lapse TIRF images were compared. The resulting coordinate-based profiles were processed using custom Python scripts. RMSD was calculated by integrating displacement magnitudes along the x- and y-axis.

### FRAP experiment

Tamoxifen-treated dynamin TKO cells were seeded on glass coverslips (CS-25R15) in a six-well plate at 1.5 × 10^5^ cells/well and transfected with Cav1-GFP as described above and incubated overnight (37°C, 5% CO_2_). Cells were serum-starved for one hour prior to treatment with DMSO (0.1%) or Dyngo-4a (30uM) for 30 min before imaging using a Zeiss Spinning Disk Confocal microscope (63X lens). Three reference images were recorded before a ROI was photobleached for 1000 ms using maximal laser intensity (488 nm and 561 nm). The fluorescent recovery images were taken every 3 s for 10 min. The signal recovery was monitored in focal plane close to the basal membrane. The intensities of the bleached regions were corrected for background signal and photobleaching of the cell. Data from at least 12 cells were collected per condition and mean FRAP recovery curves were plotted using Prism 10.0.2

### Image deconvolution and GP analysis of C-Laurdan images

The images were deconvolved to improve the resolution (57). The image stacks were first cropped around single cells. Image deconvolution was performed using Huygens Professional (SVI, Netherlands) with default settings and automatic background detection. A theoretical point spread function (PSF) was used and the maximal number of iterations was set to 50. The upper and lower channels (CH483 and CH442)) were deconvolved concurrently with a signal to noise ratio of 1. The concurrently deconvolved images were saved using a linked scale intended to retain their relative intensities. The regions of interest (ROIs) were created using the ImageJ macro Find_plasma_membrane (58), where the center images from the deconvolved z-stacks from the two channels were aligned using the ImageJ plugin *StackReg* (59). Only plasma membrane stretches without adjacent stained intracellular membranes were selected for the analysis. First, a segmented line ROI was created by manually selecting a few points close to the plasma membrane section to be analyzed. Then, the pixels with the highest intensity along lines perpendicular to each selected point were identified, additional points interpolated between them and their positions optimized in the same manner. The background was subtracted based on the mean intensities in an area outside the cells. Generalized polarization (GP) values were calculated for all the pixels in the plasma membrane ROIs according to the formula:

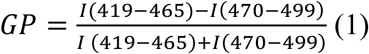

### Generation of SLBs and analysis by QCM-D

POPC vesicles for forming SLBs were prepared by diluting POPC in chloroform:methanol (3:1 volume/volume), drying it under a stream of nitrogen and rehydrating in 20 mM HEPES buffer, 150 mM NaCl (pH 7.4) (HBS buffer) at a final concentration of 1 mg/ml. Liposomes were sonicated using a bath sonicator and then extruded 11 times (Mini Extruder, Avanti) through a polycarbonate filter (Nuclepore Track-Etched Membranes, Whatman) with 100-nm pore size. An X4 unit (AWSensors) equipped with four flow chambers was used to conduct the QCM-D measurements. Wrapped 14-mm (5 MHz, Cr/Au–SiO_2_, polished) sensors were used for all experiments. Each sensor was stored in 2% SDS overnight and treated with UV/ozone (BioForce Nanosciences) for 30 min before use. The frequency and dissipation changes for overtones 1, 3, 5, 7, 9, and 11 were all recorded, but only the third overtone is reported herein. POPC vesicles (100 μL, 0.1 mg/mL) in HBS buffer were injected in a continuous flow and SLB formation was monitored. After SLB formation, the chambers were rinsed with HBS buffer. After reaching a stable baseline, DMSO (0.2%) was injected into the chamber. The flow was paused once the solution had filled the sensor chamber, and the system was allowed to reach equilibrium before 30 µM Dyngo-4a in DMSO (0.2%) or DMSO (0.2%) alone as control was injected into the chamber. The flow was paused again once the solution had filled the sensor chamber, and the system was allowed to reach equilibrium before rinsing with the buffer. Experiments were run in triplicats.

### Estimation of Dyngo-4a retained in SLBs

The adsorbed mass of Dyngo-4a was calculated from the frequency shift using the Sauerbrey Equation (60) using the AWSensors software (AWS suite). The system followed the necessary criteria for valid calculations: 1) thin adsorbed layer relative to the sensor, 2) homogenous adsorbed layer and 3) ridged adsorbed layer (61). The POPC SLBs were very thin (∼4 nm) and homogenous displaying 2D fluidity. The SLB rigidity, before DMSO addition and after DMSO removal, was established to ΔD/ΔF < 6.6×10^-8^ Hz^-1^ as measured by the AWSensors using the third overtone of a 5MHz sensor (Fig. S2). The clean sensor surface before SLB formation was used as the zero reference. The mole percentage of Dyngo-4a retained in the SLB after rinsing was calculated by using the molecular weights of POPC (759.578 g/mol) and Dyngo-4a (338.3 g/mol).

### Lipid Monolayer Experiments

Lipid monolayer experiments were performed with a Microtrough G1 System (Ø 53 × 4 mm, Kibron). The trough was covered to prevent temperature and humidity loss. POPC lipid was prepared at a total lipid concentration of 1 mM in chloroform:methanol (3:1 vol/vol). A microbalance equipped with a Wilhelmy plate was used to measure the surface pressure (π) and calibrated before each measurement. Lipid solutions were deposited onto the surface of the subphase (20 mM HEPES, 150 mM NaCl, pH 7.4) to obtain the initial surface pressure (π_0_). The subphase was continuously stirred by a magnetic stirrer. After the solvent had been allowed to evaporate for 15 min and a stable monolayer was formed, Dyngo-4a in DMSO or DMSO was injected under the lipid film directly into the subphase using a thin syringe needle (final concentration 30 μM Dyngo-4a, 0.1% DMSO). Experiments were run in triplicates with initial surface pressures between 20-23 mN m^-1^ prior to injection of Dyngo-4a yielding similar results and analysis of the adsorption curves was performed with GraphPad Prism.

### MD simulations

The CHARMM force field (62–64) was used to run MD simulations. The Dyngo-4a molecule was built using ligand modeler (65) and validated by octanol-water partitioning compared to cLogP value provided by the supplier (cLogP = 2.6; calculated from simulations = 2.5). Two membrane compositions were studied: pure POPC, and a POPC:cholesterol mixture at a 70:30 molar ratio. Each POPC:CHOL system comprised a fully hydrated bilayer of 70 POPC and 30 cholesterol molecules (∼10,000 water molecules), while the POPC-only system contained 76 POPC lipids. Initial equilibration was performed on Dyngo-4a–free membranes. Subsequently, Dyngo-4a were added randomly in the aqueous phase, either as single molecules or in sets of ten, followed by additional energy minimization and equilibration. Simulation box dimensions were approximately 5 × 5 × 16 nm³, sufficient to minimize finite-size artifacts and maintain the minimum image convention. Periodic boundary conditions were applied in all directions. All simulation series were run in triplicates and presented as average and standard deviation. Initial equilibration simulations were conducted using the Nose-Hoover thermostat (66) (and semi-isotropic Parrinello-Rahman (67) barostat (were employed to control temperature and pressure in the simulations. For AWH simulations, a convolved adaptive biasing potential (68) was applied with a force constant of 12800kJ mol^-1^ nm^-2^ defining the curvature of the biasing potential along the reaction coordinate. Verlet cut-off (69) scheme was applied with a cut-off radius of 1.2 nm set to both van der Waals and electrostatic interactions. LINCS algorithm (70) was used to constraint all bonds with hydrogen atoms. Area per lipid was calculated by dividing the box area in the xy-plane by the number of lipids per leaflet, using output from gmx box. Deuterium order parameters were computed with gmx order, treating each leaflet separately to capture asymmetric effects. Mass density profiles along the membrane normal (z-axis) were obtained using gmx density. For the colocalization analysis two-dimensional heatmaps of lipid and Dyngo-4a distributions in the membrane plane were generated with gmx densmap, and visualized using xpm2ps and custom Python scripts to produce contour plots. All analyses were performed over the final 100 ns of each trajectory and averaged over triplicates unless stated otherwise.

### Electron Microscopy

Electron microscopy was performed as described previously (71). Briefly, cells were initially fixed in 2.5% glutaraldehyde, then underwent a series of post-fixative and contrasting immersions. These were 1.5% potassium ferricyanide and 2% osmium tetroxide in ddH_2_O, 1% thiocarbohydrazide in ddH_2_O, 2% osmium tetroxide in ddH_2_O, lead aspartate (20mM lead nitrate, 30mM aspartic acid, pH 5.5), and 1% uranyl acetate in ddH_2_O. Each step was irradiated for 3min in a microwave (BioWave) at 80W for 3min, with thorough ddH_2_O washes between steps. Cells were then serially dehydrated with ethanol before serial infiltration with LX112 resin (Ladd) and polymerisationovernight in an oven set at 60°C. Sections were acquired using a UC6 ultramicrotome (Leica) and ultrathin sections imaged using a JEM-1400 transmission electron microscope operating at 80kV.

### AFM experiments

HeLa cells were seeded on a 35 mm glass bottom Petri dish (Cellvis, Cat. # D35-10-1.5-N) and starved as described above. The cell media was exchanged to serum-free media with 20 mM HEPES containing either Dyngo-4a (30 μM) or DMSO (0.1%) and set for 30 minutes before performing AFM experiments. All AFM experiments were conducted by NanoWizard 4 XP bioscience system (Bruker/JPK, Berlin, Germany) at ambient temperature. Commercial colloid probes (Si_3_N_4_ AFM probes, 0.03 N/m cantilever; 4.5 µm polystyrene particle from Novascan Technologies, Ames, IA USA) were used to collect force maps on Cav1 FlpIn T-REx HeLa cells, with and without the treatment of Dyngo-4a after calibrating the cantilever. The spring constant for each cantilever was calculated based on the thermal noise method. A glass dish with cells was mounted on an inverted Nikon Eclipse Ti-E2 microscope (Nikon Corp., Japan) with a Prime 95B sCMOS camera (Photometrics Teledyne, USA) to localize cells using a 40x objective. The cantilever was carefully centred on cells. The whole duration of experiments was kept below 2 hours. For both conditions, force maps on 8 or 9 individual cells were collected. An approach curve with a length of 5 µm at a speed of 2 µm/s and a retract curve with a length of 7 µm and a speed of 4 µm/s at the maximum force load of 1 nN was used for mapping 8×8 matrix (or 64 points) over 10 µm^2^ scan size on individual cells and glass (i.e., a region of a glass dish with no cells). All experiments were carried out independently at least twice, with separately but identically prepared cells and AFM tips.

### AFM data analysis

Individual force curves from force maps were analyzed using the JPK data processing software. Before fitting the force curves, data was smoothed with a Gaussian filter with a width of 1. After smoothing and setting the measured force close to the contact point and offset to zero of the extend curve, the Young’s modulus (*E*) was determined by fitting with the built-in Hertz/Sneddon model (Eq. 1). The major assumption of this model is that the indentation depth (∼0.5 to 2 µm) is very small or nearly negligible as compared to the thickness of the cells (10 to 12 µm). This model also assumes that the sample is elastic and homogenous. As this is not the case for cells, we fitted the data from 0 to 500 nm of indentation (*δ*) from a contact point for both conditions.

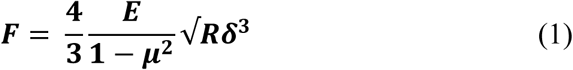

where *F* is force, R represents the radius of the spherical probe (2.25 µm) and *µ* is the Poisson ratio. The ratio varies from 0 to 0.5 depending on the material properties. For simplicity, we set it to 0.5. Curves with the root mean square (RMS) value below 15 pN were discarded from further analysis.

## Statistical analysis

Two-sample *t* test was used to compare values in the different experiments using the GraphPad prism program version 10.0.2. P < 0.05 was considered a statistically significant change. *P < 0.05; **P < 0.01; ***P < 0.001; NS, not significant. All the values were presented as mean ± standard deviation (SD) as specified in the figure legends.

## Data sharing plans

All data needed to evaluate the conclusions in this article are included in the article and/or SI appendix.

## Online supplementary material

Fig. S1 show the molecular structures of Dyngo-4a and cholesterol, immunofluorescent stainings of cavin1 and EHD2 in Dynamin TKO cells and an illustration of caveola mobility and TIRF imaging. Fig. S2 show representative images of CTxB internalization in cells treated with Dyngo-4a as well as confocal FRAP time-lapse series and quantification of repletion of cytosolic Cav1-GFP in a photobleached area in ctrl or Dyngo-4a treated KO cells. Fig. S3 show the softness plot from QCM-D measurements during SLB formation. Fig. S4 provides additional MD simulation data of ten Dyngo-4A molecules in POPC bilayer or in POPC:Chol bilayer. Fig. S5 shows the analysis of the effect of Dyngo-4a on PM mobility, caveolae displacement and Tfn uptake as well as AFM force-indentation curves, and incorporation of Chol using fusogenic liposomes and MβCD. Video 1 show TIRF microscopy timelapse of a Dynamin TKO cell treated with DMSO (0.1%) for 30 min. Video 2 show TIRF microscopy timelapse of a Dynamin TKO cell treated with Dyngo-4a (30µM) for 30 min. Video 3 shows the association of cavin1 and Cav1-GFP following Dyngo-4a-treatment.

