## Supplementary Information for "Lipid packing contributes to the confinement of caveolae to the plasma membrane"

**Affiliations**

**This PDF file includes:**

Figs. S1 to S5 and legends to Video 1 – 3.

### Supplemental Figure legends

#### Figure S1

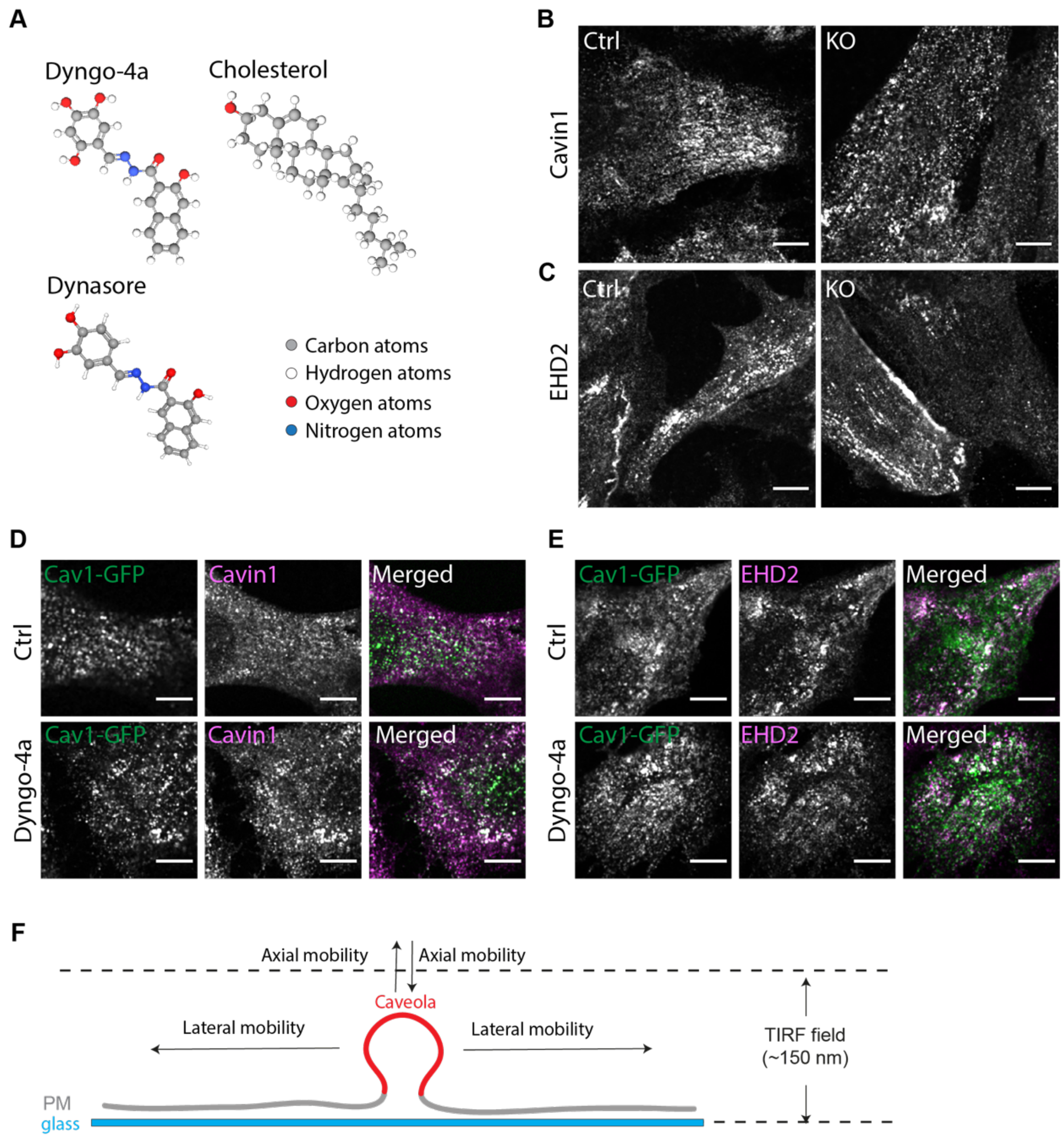


**Figure S1. Cavin1 and EHD2 expression in dynamin triple knock out fibroblasts.** Ball-and-stick representations of Dyngo-4a (top left panel), Dynasore (bottom left panel) and Cholesterol (top right panel). Carbon atoms are shown in grey, hydrogen in white, oxygen in red, and nitrogen in blue. **(B-C)** Representative immunofluorescent staining of dynamin triple knock out cells showing Cavin1 (B) and EHD2 (C) expression after 5 days of tamoxifen-treatment (KO) or no treatment (Ctrl) as indicated. Scale bar, 5 µm. **(D-E)** Representative immunofluorescent staining of Cav1-GFP expressing KO cells showing Cavin1 (D) and EHD2 (E) expression after 30 min treatment with (0.1%) DMSO (top panels) or 30 µM

Dyngo-4a (bottom panels). Scale bar, 10 µm. **(F)** Illustration of caveola mobility and detection in TIRF imaging. **(G)** Representative confocal FRAP time-lapse series showing repletion of cytosolic Cav1-GFP in a photobleached area in ctrl or Dyngo-4a treated KO cells as indicated. Intensity recovery was monitored for 10 min after photobleaching. Cav1-GFP fluorescence intensities is intensity-coded using LUT. Scale bar, 5 µm. (**H**) Recovery curves of Cav1-GFP corresponding to (G). Cav1-GFP fluorescence intensity was normalized to background and reference. $n$ =12, mean ± SEM.

### Figure S2

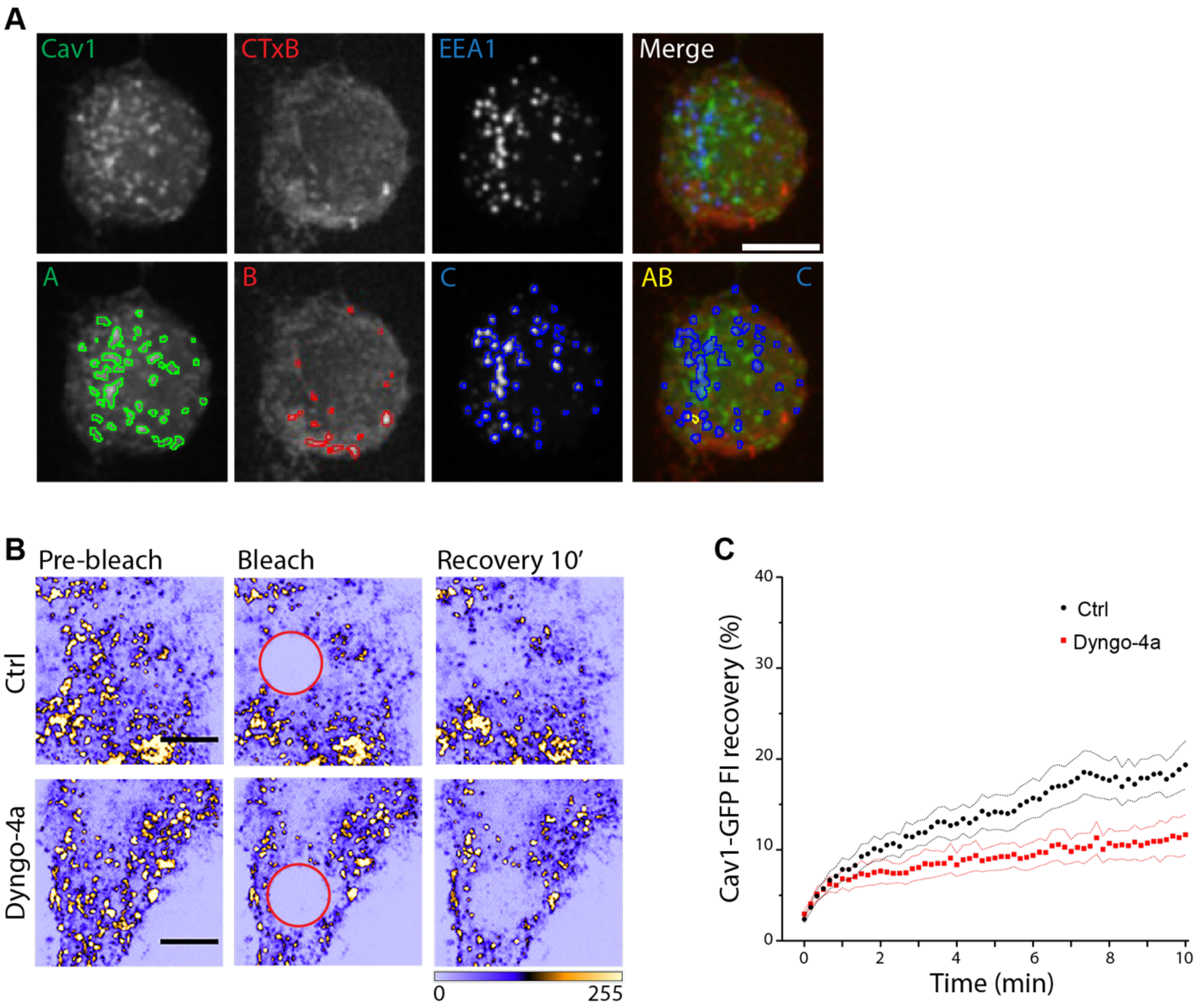


**Figure S2. Caveolae internalization is inhibited by Dyngo-4a. (A)** Representative immunofluorescent maximum-intensity projections of fluorescent CTxB uptake in Dyngo-4a-treated cells. Bottom panels show the masks generated from each fluorescent marker which were used to quantify the number of Cav1-GFP and CTxB-647 colocalizing spots devoid of EEA1. **(B)** Representative confocal FRAP time-lapse series showing repletion of cytosolic Cav1-GFP in a photobleached area in ctrl or Dyngo-4a treated KO cells as indicated.

Intensity recovery was monitored for 10 min after photobleaching. Cav1-GFP fluorescence intensities is intensity-coded using LUT. Scale bar, 5 μm. (**C**) Recovery curves of Cav1-GFP corresponding to (G). Cav1-GFP fluorescence intensity was normalized to background and reference. *n* =12, mean ± SEM.

**Figure 3**

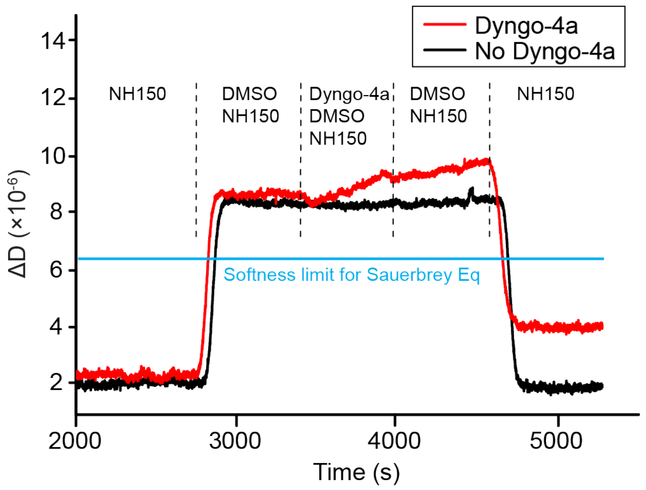


**Figure S3. Dyngo-4a adsorbs to and affects the softness of the SLB.** QCM-D measurements of the softness (ΔD) during SLB formation and Dyngo-4a (30 μM) (red line) or DMSO (0.2%) (black line) adsorption or desorption over time. Arrows depict time of injection of DMSO (0.2%, equivalent to the concentration that Dyngo-4a is dissolved in), sample injection (Dyngo-4a or DMSO), 0.2 % DMSO injection and saline buffer wash (150 mM NaCl, pH 7.4) injection respectively.

**Figure S4**

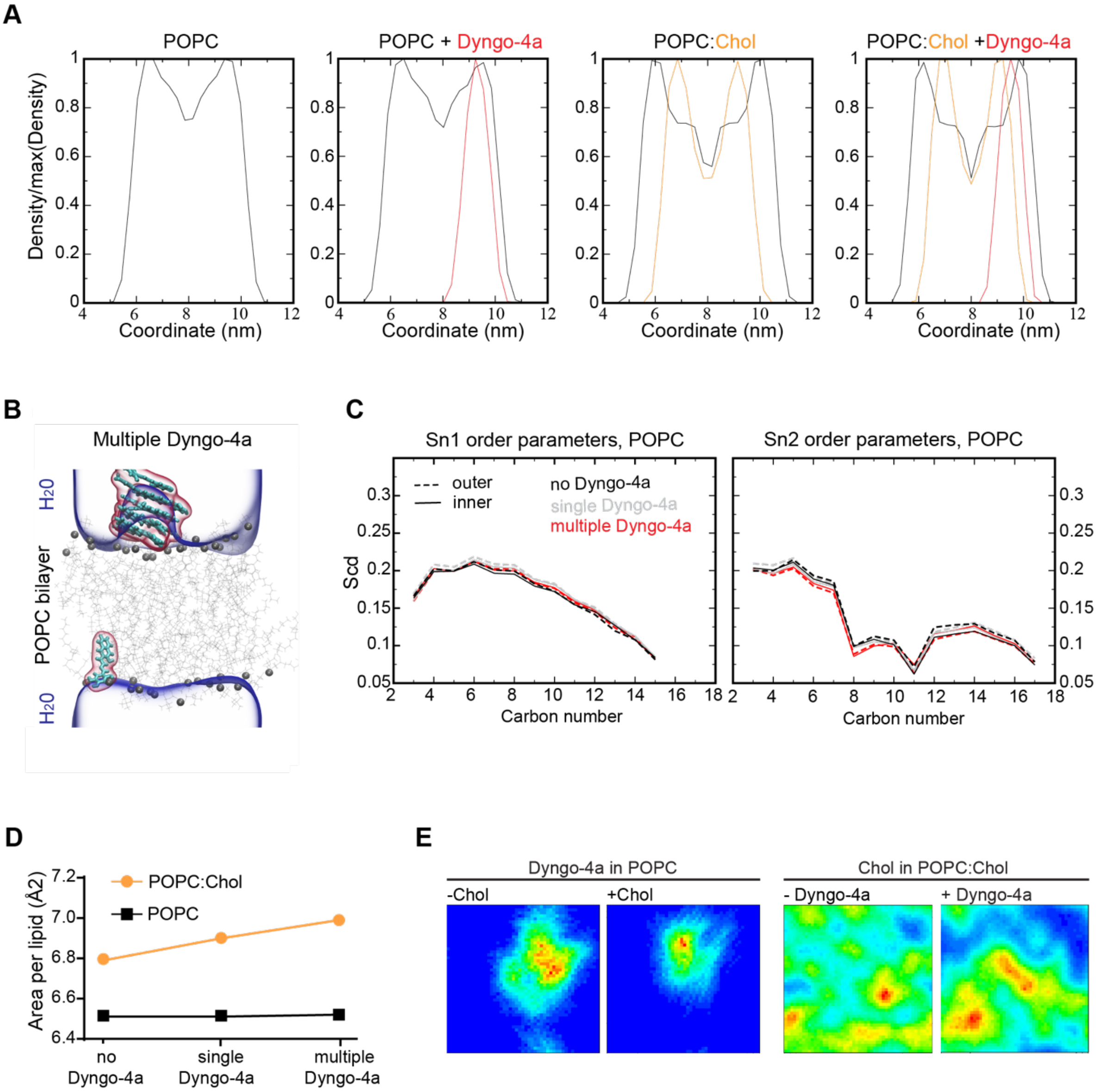


**Figure S4. Dyngo-4a triggers lipid packing frustration in POPC:Chol membranes.** **(A)** Normalized density profiles for POPC (black), POPC:Chol (30:70, orange) membranes without and with inserted Dyngo-4a (red) molecules. The center of the membrane is located at 8 nm on the x axis of the graphs. **(B)** Snapshot of multiple Dyngo-4a molecules (cyan stick and red volume) in POPC bilayer. Dyngo-4a molecules did not cross the membrane throughout the simulation so they were placed randomly at the beginning of the simulations and could translocate from one leaflet to another via periodic boundary condition. Phospholipid headgroups shown in dark gray and lipid chains as light gray sticks, dark blue represents the aqueous phase. **(C)** Deuterium order parameter profiles for POPC tails in a POPC membrane.

Left show sn1 tail (saturated, 16 carbons) and right shows sn2 tail (unsaturated, 18 carbons). No Dyngo-4a (black), single Dyngo-4a (gray), multiple Dyngo-4a (red), dashed lines depict the outer leaflet and solid line the inner leaflet. **(D)** Plot showing how the area per lipid changes upon addition of a single or multiple Dyngo-4a molecules in a POPC (black) or POPC:Chol (70:30, orange) membrane. **(E)** Heatmaps depict the spatial distribution of Dyngo-4a in POPC or POPC:Chol (70:30) membrane (left) or the spatial distribution of Chol in POPC:Chol (70:30) membrane with or without Dyngo-4a (right) as indicated. Dyngo-4a displays higher mobility in a pure POPC membrane compared to when it neighbors a chol cluster in a POPC:Chol membrane. Chol display less uniform distribution in the membrane in the presence of Dyngo-4a.

**Figure S5**

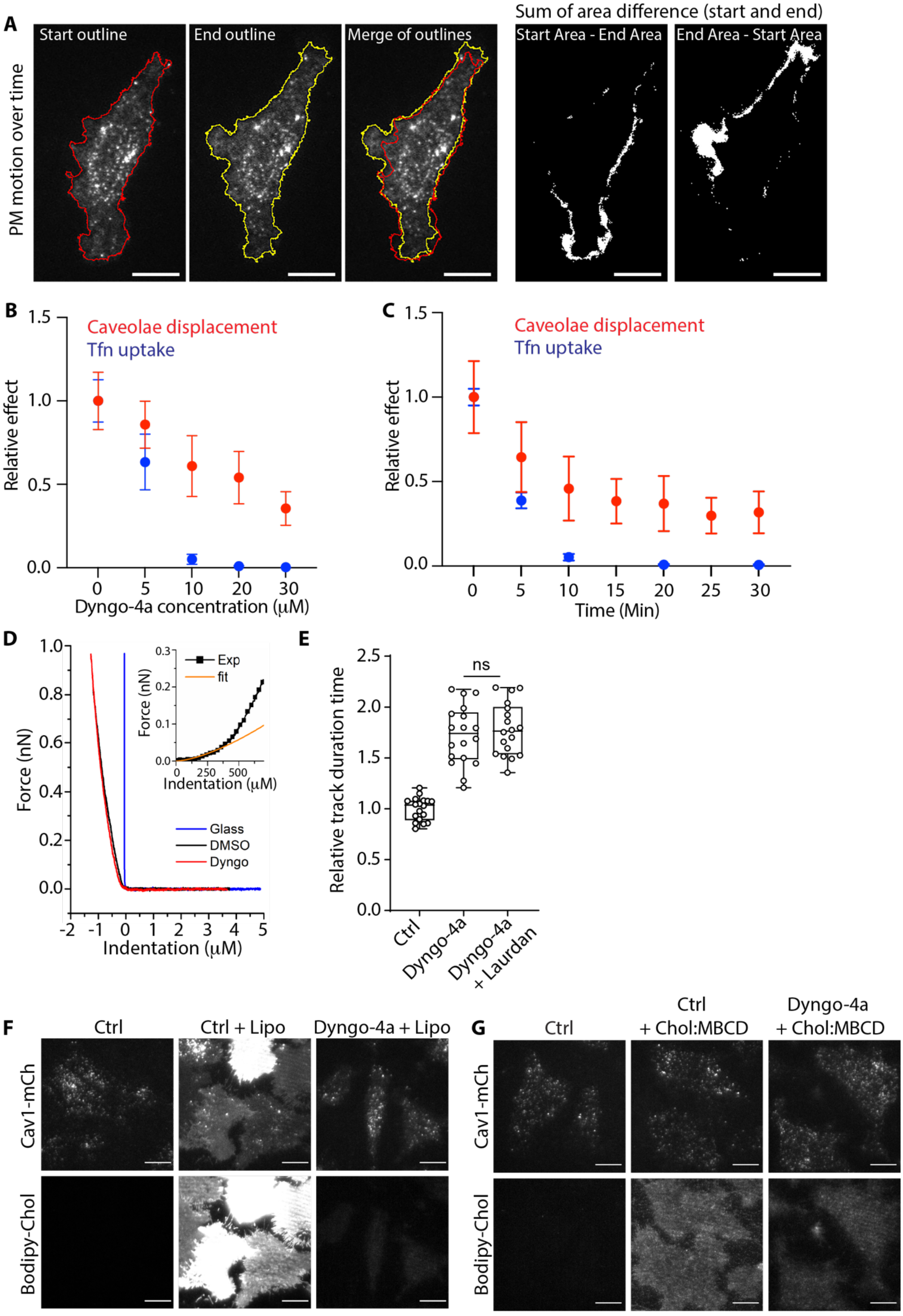
A
PM motion over time
Start outline
End outline
Merge of outlines
Sum of area difference (start and end)
Start Area - End Area
End Area - Start Area
B
Caveolae displacement
Tfn uptake
Relative effect
Dyngo-4a concentration (μM)
C
Caveolae displacement
Tfn uptake
Relative effect
Time (Min)
D
Force (nN)
Indentation (μM)
Exp
fit
Glass
DMSO
Dyngo
E
Relative track duration time
ns
Ctrl
Dyngo-4a
Dyngo-4a + Laurdan
F
Ctrl
Ctrl + Lipo
Dyngo-4a + Lipo
Cav1-mCh
Bodipy-Chol
G
Ctrl
Ctrl + Chol:MBCD
Dyngo-4a + Chol:MBCD
Cav1-mCh
Bodipy-Chol

**Figure S5. Analysis of the effect of Dyngo-4a on PM mobility, caveolae displacement and Tfn uptake, AFM force-indentation curves, and incorporation of Chol using fusogenic liposomes and MβCD. (A)** Representative TIRF micrographs and illustration of how the PM motion was retained from a 5 minutes TIRF movie. Basal plasma membrane outline at start (red line) and at the end (yellow line). Black and white images illustrate the area change from the start image to the end image. **(B)** Plot showing calculated RMSD values preformed on plasma membrane profile sections from ctrl and Dyngo-4 treaded cells. Start and end frame of 5 min time-lapse TIRF images were compared. **(C-D)** Quantification of the caveolae displacement length (red) and Tfn-647 uptake (blue) in cells treated with DMSO (ctrl) or Dyngo-4a at different concentrations (C) and different time points (D) as indicated. Analysis of caveolae mobility was performed using Imaris software, and track means from at least 25 cells per condition are shown ± SD. Quantification of Tfn-647 intensity was performed using ImageJ and relative intensity means from at least 500 cells are shown ± SD. **(E)** Representative AFM force curves detected on glass, on DMSO-treated cells (ctrl), and Dyngo-4a-treated cells. Inset show example fitting of the force curve with Hertz model. **(F)** Quantification of Cav1-GFP track duration time in cells treated with DMSO (ctrl) or Dyngo-4a in the presence and absence of C-Laurdan. Numbers were related to ctrl-treated cells. Analysis was performed using Imaris software, and track mean from at least 30 cells per condition are shown ± SD. Significance was assessed using *t* test, **** $p \leq 0.0001$. **(G)** Representative TIRF micrographs of ctrl or Dyngo-4a treated Cav1-mCh cells 5 minutes after addition of fusogenic liposomes containing Bodipy-Chol or no addition as indicated. **(H)** Representative TIRF micrographs of ctrl or Dyngo-4a treated Cav1-mCh cells 5 minutes after addition of Bodipy-chol:MBCD or no addition as indicated. All scale bars, 10 µm.

**Video 1. DMSO-treatment does not stall caveolae mobility and plasma membrane undulations.** Representative live cell video of a Dynamin TKO cell transiently expressing Cav1-GFP, treated with DMSO (0.1%) 30 min prior to imaging. Cells were imaged on TIRF every third s for 5 min. Frame rate: 10 frames per second. Scale, 10 µm.

**Video 2. Dyngo-4a-treatment results in loss of caveolae mobility and plasma membrane undulations.** Representative live cell video of a Dynamin TKO cell transiently expressing Cav1-GFP, treated with Dyngo-4a (30µM) 30 min prior to imaging. Cells were imaged on TIRF every third s for 5 min. Frame rate: 10 frames per second. Scale, 10 µm.

**Video 3. Cavin1 remains associated with Cav1-GFP in the plasma membrane after 30 min Dyngo-4a-treatment.** Representative live cell video of a Dynamin TKO cell transiently expressing Cav1-GFP (green) and mCh-Cavin1 (magenta), treated with Dyngo-4a (30µM) 30 min prior to imaging. Cells were imaged on TIRF every third s for 5 min. Frame rate: 10 frames per second. Scale, 5 µm.
